# Uncoupling Osteoclast Fusion and Resorptive Activity Through Differential Talin1-Rap1 Signalling Pathways

**DOI:** 10.64898/2026.11.11.765932

**Authors:** Marianna Plozza, Rosa Nicole Freiberger, Arnaud Métais, Thibaut Sanchez, Alicia Deleris, Camille Gorlt, Baptiste Casel, Lotfi Slimani, Ophélie Dufrançais, Serena Giuntini, Isabella da Silva, Maxime Kaiser, Yoann Rombouts, Anne Gonzalez de Peredo, Frédérique Fallone, Renaud Poincloux, Brigitte Raynaud-Messina, Jérôme Bouchet, Christel Vérollet, Frédéric Lagarrigue

**Affiliations:** Institut de Pharmacologie et Biologie Structurale (IPBS), Université de Toulouse, Centre National de la Recherche Scientifique, Université Toulouse III, CNRS, IPBS, Toulouse, France; URP 2496, Laboratory Orofacial Pathologies, Imaging and Biotherapies, Life Imaging Platform (PIV), University Paris Cité, 1 rue Maurice Arnoux, 92120 Montrouge, France

## Abstract

Bone remodelling is essential for skeletal growth and repair and depends on a balance between osteoblast-driven bone formation and osteoclast-mediated bone resorption. Integrin activation, regulated by Rap1 and Talin1, is critical for osteoclast function by coordinated adhesion and actin cytoskeleton organization. In this study, we investigated Rap1 effectors controlling Talin1 in osteoclasts. RIAM and Lamellipodin (Lpd) are expressed and localise to actin-rich structures, including tunneling nanotubes (TNTs) and podosomes, supporting roles in cytoskeletal organisation and intercellular connectivity. Functional analyses revealed distinct, stage-specific functions. Lpd deficiency impaired early osteoclastogenesis by reducing TNT formation, disrupting precursor fusion, decreasing multinucleation, and lowering bone resorption. In contrast, RIAM deficiency has minimal impact on fusion but significantly reduced resorption activity in mature osteoclasts, indicating a role at later stages. Disruption of the direct Rap1– Talin1 interaction caused broader defects affecting both early and late stages, including reduced TNT formation, delayed fusion, altered multinucleation, and impaired bone degradation. In vivo, only disruption of this direct interaction increased bone mass and reduced osteoclast number and/or activity, whereas RIAM or Lpd deletion had no significantly impact. These results demonstrate that RIAM and Lpd perform distinct but complementary functions and identify the direct Rap1–Talin1 interaction as a central mechanism controlling osteoclast fusion and bone resorption.

## Introduction

Bones undergo continuous remodelling, a process essential for skeletal development and growth, fracture repair, and calcium homeostasis. This remodelling depends on a tightly regulated balance between osteoblasts, which form bone, and osteoclasts, which resorb it (Blangy *et al*., 2020). Osteoclasts are multinucleated giant cells generated through the fusion of mononuclear haematopoietic precursors from the monocyte/macrophage lineage, under the control of receptor activator of nuclear factor kappa-B ligand (RANKL) and macrophage colony-stimulating factor (M-CSF) (Boyle, Simonet and Lacey, 2003). As with other myeloid cells, mature osteoclasts display considerable heterogeneity: they are highly efficient at fusion, but can also undergo fission and recycle into new osteoclasts (Madel *et al*., 2019; McDonald *et al*., 2021). The formation of multinucleated osteoclasts is a key step, as the degree of multinucleation most often correlates with bone-resorbing activity (Dufrançais *et al*., 2021). This fusion process is regulated in vivo at homeostasis (Barnea-Zohar *et al*., 2021), and enhanced in pathological conditions such as bone inflammation, certain infections (e.g. HIV-1), cancers and age-related osteoporosis, leading to highly degradative osteoclasts. Their central role in bone disease highlights the need to better understand the mechanisms regulating osteoclast formation and function.

Each stage of osteoclast differentiation, known as osteoclastogenesis, involves complex cellular processes driven by actin cytoskeleton remodelling and integrin-mediated adhesion and fusion (Dufrançais *et al*., 2021). We recently showed that in the early stages, fusion between mononuclear precursors is facilitated by the formation of tunnelling nanotubes (TNTs), intercellular bridges containing F-actin and microtubules, known to direct intercellular communication (Dufrançais *et al*., 2021, 2025; Zurzolo, 2021). In later stages, osteoclast function involves the formation of actin-based adhesive structures called sealing zones, consisting of a complex and dense assembly of podosomes (Jurdic *et al*., 2006; Portes *et al*., 2022). Podosomes are highly dynamic F-actin structures that mediate adhesion via integrins, sense mechanical cues, and degrade extracellular matrix, thereby contributing to tissue remodelling and cell invasion. These structures are specific to cells of the monocyte lineage, such as macrophages and osteoclasts, and consist of a central F-actin core surrounded by a ring of adhesion proteins, notably integrins. In mature osteoclasts, the assembly of podosomes into larger superstructures, i.e., the sealing zone, is absolutely essential for efficient bone resorption (Jurdic *et al*., 2006; Teitelbaum, 2011; Blangy *et al*., 2020). However, how these actin-based structures and their molecular components adapt to different physiological or pathological contexts remains unclear.

Integrins are heterodimeric αβ receptors expressed at the cell surface that mediate interactions with extracellular matrix proteins and other cells (Hynes, 2002). Osteoclasts express several integrin families, including β1, β2, and β3, all of which are essential for bone resorption (Schmidt *et al*., 2011). Among them, the presence of integrin αvβ3 in the sealing zone of mature osteoclasts plays a predominant role in adhesion and is widely used as a marker of their differentiation (Teitelbaum, 2011). A key property of integrins is their ability to switch between low- and high-affinity states for their ligands. This activation process is driven by the binding of Talin1 and Kindlin3 to the cytoplasmic domains of β1, β2, and β3 integrins (Shattil, Kim and Ginsberg, 2010; Calderwood, Campbell and Critchley, 2013). Talin1 is a large cytoskeletal adaptor composed of an N-terminal FERM head domain (subdivided into F0–F3 subdomains) and an extended C-terminal rod domain containing multiple helical bundles (R1-R13) and binding sites for actin and regulatory proteins. Together, Talin1 and Kindlin3 link integrins to the actin cytoskeleton and transmit signals from the extracellular environment (Calderwood, Campbell and Critchley, 2013). In the absence of Kindlin3, osteoclasts form only immature podosomes, cannot establish sealing zones, and exhibit severely impaired bone resorption, leading to osteopetrosis (Schmidt *et al*., 2011). Similarly, Talin1-deficient osteoclast precursors exhibit defective adhesion, migration, and cytoskeletal organization, resulting in an inability to mature into functional resorbing cells, leading to increased bone mass in mice (Zou *et al*., 2013). Rap1, a small GTPase of the Ras family, plays a central role by promoting the recruitment of Talin1 to the plasma membrane and facilitating its interaction with integrins (Han *et al*., 2006). Loss of Rap1 in mature osteoclasts blocks bone resorption and increases bone mass (Zou *et al*., 2013). Moreover, selective inhibition of Rap1 or Talin1 in osteoclasts protects against pathological bone loss, such as that induced by ovariectomy or inflammatory arthritis (Zou *et al*., 2013). However, the effector(s) of Rap1 responsible for the regulation of Talin1 in osteoclasts remains unknown.

A yeast two-hybrid screen for Rap1-GTP-interacting proteins in T lymphocytes identified RIAM (Rap1-GTP-interacting adaptor molecule), a key mediator of Rap1-dependent integrin activation and actin regulation (Lafuente *et al*., 2004). In parallel, a ligand screen for Ena/VASP proteins identified Lamellipodin (Lpd) as the murine orthologue of human RIAM (Krause *et al*., 2004). Both RIAM and Lpd belong to the MRL (MIG-10, RIAM and Lamellipodin) family of adaptor proteins, which regulate actin dynamics, cell adhesion and migration, and cell growth. These proteins share a conserved core region containing a Ras-association domain and a pleckstrin homology domain (Krause *et al*., 2004; Lafuente *et al*., 2004), while their N-terminal regions bind to Talin1 within its rod domain, primarily at the R2–R3 helical bundle region (often referred to as the RIAM-binding site), thereby promoting Talin1 recruitment to the plasma membrane (Lee *et al*., 2009; Calderwood, Campbell and Critchley, 2013). Their C-terminal regions are more variable and interact with actin-regulatory proteins such as Ena/VASP, profilin, and Abi/WASP (Krause *et al*., 2004; Lafuente *et al*., 2004; Law *et al*., 2013). MRL proteins localise both in the cytoplasm and in lamellipodia. RIAM binds primarily to Rap1, and to a lesser extent to active Ras (Lafuente *et al*., 2004; Han *et al*., 2006; Lee *et al*., 2009), whereas Lpd appears to preferentially interact with other GTPases such as Rac1 and M-Ras (Tasaka, Negishi and Oinuma, 2012; Law *et al*., 2013), suggesting distinct signalling roles. Functionally, RIAM is highly expressed in leukocytes and regulates key immune processes, including β2 integrin activation, leukocyte recruitment, immunological synapse formation, and complement-dependent phagocytosis (Klapproth *et al*., 2015; Su *et al*., 2015; Lagarrigue *et al*., 2017; Sun *et al*., 2021). In contrast, Lpd has a broader role in cell migration and, within the immune system, is mainly involved in regulatory T cells, where it supports integrin function and suppressive activity (Law *et al*., 2013; Carmona *et al*., 2016; Dimchev *et al*., 2020; Sun *et al*., 2021). Consistent with these roles, RIAM deficiency protects mice from T cell-mediated autoimmune diseases such as colitis and type 1 diabetes (Lagarrigue *et al*., 2017; Sun *et al*., 2021).

In addition to indirect regulation via RIAM and Lpd, Rap1 can bind directly to Talin1. Structural studies have identified two Rap1-binding sites within the F0 and F1 subdomains of Talin1 (Zhu *et al*., 2017; Lagarrigue *et al*., 2018; Gingras *et al*., 2019). These interactions are essential for integrin function in platelets and for haemostasis (Lagarrigue *et al*., 2020). Disrupting Rap1–Talin1 binding leads to measurable defects in the activation of several integrins (αLβ2, α4β1, α4β7) in CD4⁺ T cells (Lagarrigue *et al*., 2022). These defects can be partially rescued by RIAM overexpression or worsened by its deletion, highlighting cooperation between pathways (Lagarrigue *et al*., 2022). Similarly, Rap1–Talin1 interactions, together with Lpd, contribute to integrin regulation in regulatory T cells (Lagarrigue *et al*., 2022). Overall, these findings support a model in which multiple Rap1-dependent pathways converge on Talin1 to control integrin activation.

In this study, we examined how the three Rap1-dependent effector pathways that regulate Talin1 contribute to osteoclast differentiation and bone resorption activity. Using mouse osteoclasts genetically modified for RIAM or Lpd and a unique model of disruption of the direct Rap1–Talin1 interaction, we found that RIAM and Lpd have distinct but complementary functions and identified the direct Rap1–Talin1 interaction as a key mechanism controlling osteoclast fusion and bone resorption.

## Results

### RIAM and Lpd localise to actin-based adhesive structures in osteoclasts

The osteoclast differentiation process is characterized by a series of highly coordinated cellular events, primarily governed by the dynamic remodelling of the actin cytoskeleton, associated with integrin-mediated adhesion and membrane fusion (Dufrançais *et al*., 2021). We performed immunostaining to visualize Talin1, RIAM, and Lpd proteins in murine bone marrow-derived osteoclasts (Figure 1) and in mature human osteoclasts differentiated from circulating monocytes (Figure S1). Interestingly, the spatial distribution of these proteins reveals their presence or accumulation on TNTs (Figure 1, left), intercellular bridges composed of filamentous actin (F-actin) that do not touch the substrate. These structures are known to facilitate cytoplasmic continuity and it has been described that they participate in cell fusion between osteoclast precursors (Dufrançais *et al*., 2025). In addition, Talin1 exhibits marked enrichment around the actin core of podosomes within the podosomal belt, a specialized adhesive structure surrounding the cell periphery and composed of multiple podosomes (Figure 1A and S1). This localization is consistent with the established role of Talin1 in the binding of integrin to the actin cytoskeleton and the regulation of adhesion dynamics, as previously described in osteoclasts (Jurdic *et al*., 2006; Teitelbaum, 2011; Portes *et al*., 2022). Similarly, RIAM is highly expressed within podosome belts (Figure 1B and S1), particularly around the F-actin cores of podosomes, which is consistent with previous observations in macrophages (Klapproth *et al*., 2015). Although Lpd is also detected at podosomes, its localization profile is more heterogeneous; Lpd is also enriched at intercellular contact sites in mouse and human osteoclasts (Figure 1C and S1), suggesting a potential role in coordinating membrane interaction and cytoskeletal organization during cell fusion. Collectively, these observations demonstrate that, like Talin1, RIAM and Lpd are expressed in osteoclasts and are specifically recruited to actin-based structures.

**Figure 1:**
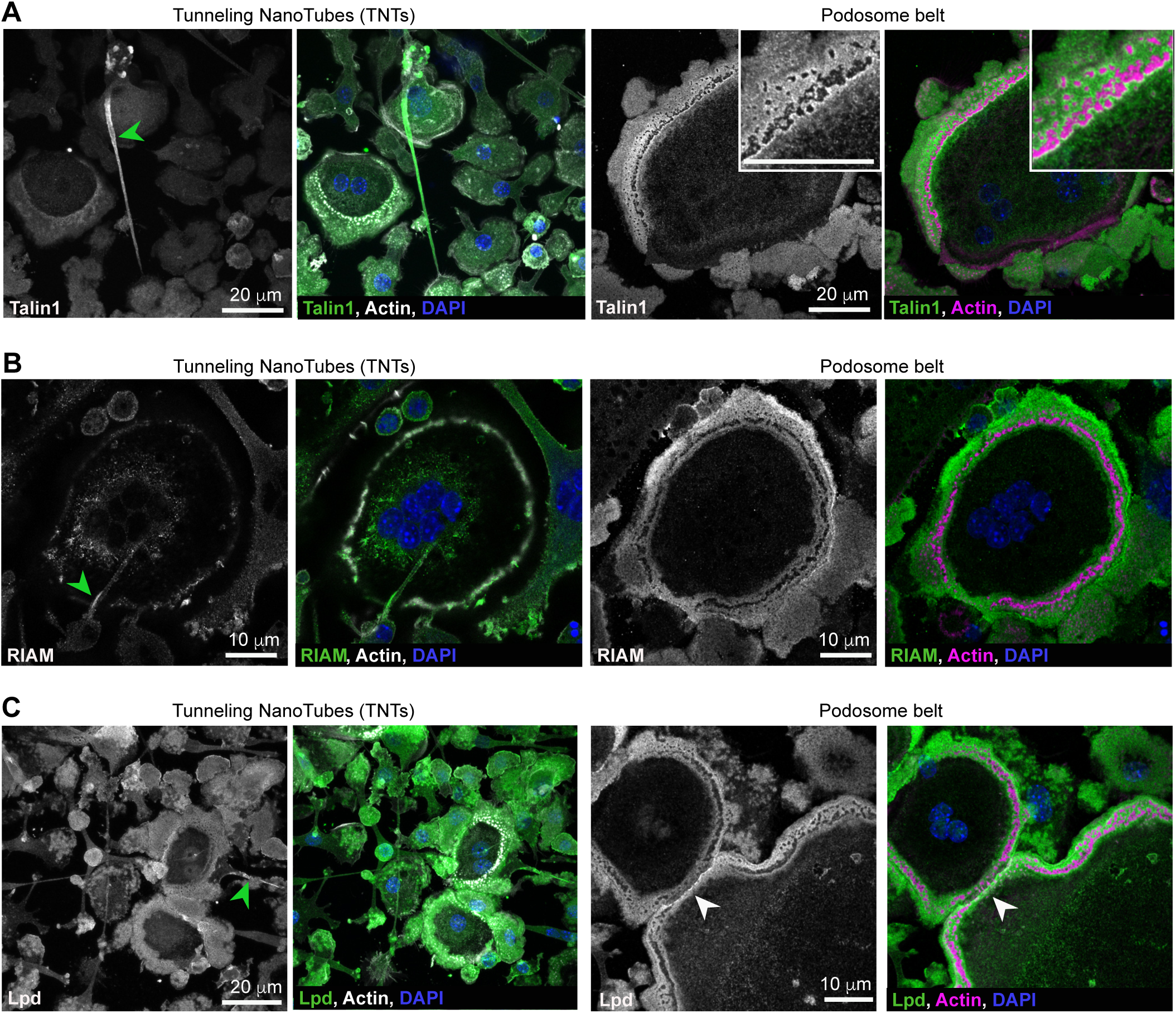
Talin1, RIAM and Lpd localize at F-actin structures in murine osteoclasts. Confocal microscopy images showing the localization of Talin1 **(A)**, RIAM **(B)** and Lpd **(C)** in mouse osteoclasts cultured on glass slides at day 4 of differentiation: F-actin (phalloidin, white on the left and magenta on the right), nuclei (DAPI, blue), and Talin1/RIAM/Lpd (green). F-actin structures: tunneling nanotubes (TNTs, left) and podosome belts (right). Scale bars: 10 or 20 µm. Green arrowheads indicate accumulation on TNTs, and white arrowheads indicate Lpd accumulation at cell-contact sites.

### Differential contributions of RIAM and Lpd to osteoclast fusion and multinucleation

The preferential localisation of RIAM and Lpd within structures implicated in osteoclast formation through precursor fusion prompted us to evaluate their functional contribution to this process, and to compare it to that of Talin1. To do this, we used a murine model of myeloid progenitor differentiation based on conditional immortalisation via the homeobox gene HoxB8. These progenitors were then differentiated into mature osteoclasts by stimulation with M-CSF and RANKL, thus reproducing in vitro the different stages of osteoclastogenesis, as previously described (Dufrançais *et al*., 2025).

To generate near-pure osteoclast populations deficient in Talin1, RIAM, or Lpd, we established immortalised myeloid progenitors derived from mice carrying floxed alleles of each respective gene. These progenitors were transduced with a lentiviral vector expressing Cre recombinase fused to the fluorescent marker mCherry. This strategy enabled the isolation, by flow cytometry, of cells positive for both mCherry and eYFP, the latter serving as a reporter of Cre recombinase activity (Figure S2). Control cells transduced with a lentivirus expressing only mCherry did not express eYFP, as expected (data not shown). Flow cytometry analyses confirmed that osteoclasts derived from these populations were indeed lacking the target proteins (Figure 2A).

**Figure 2:**
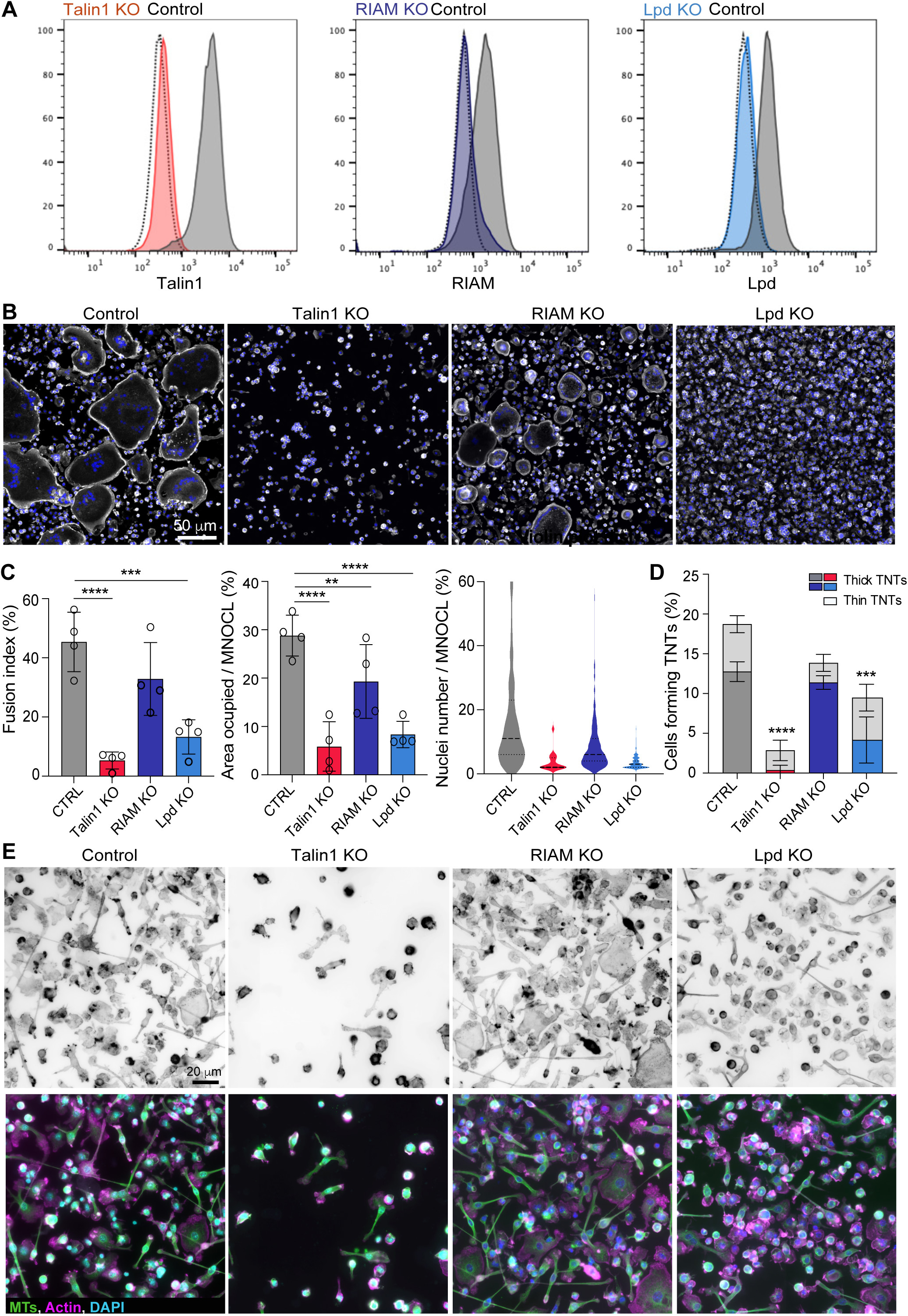
Distinct role of RIAM and Lpd on tunneling nanotube (TNT) formation and osteoclast fusion. **(A)** Flow cytometry analysis of Talin1, RIAM, and Lpd expression in mature murine Talin1 KO, RIAM KO, Lpd KO, and control osteoclasts. Representative histograms (n=3 experiments). **(B-C)** Cell fusion analysis. Microscopy analysis of cell fusion in control and Talin1, RIAM or Lpd KO osteoclasts on day 4 of differentiation. **(B)** Representative microscopy images: F-actin (phalloidin, white) and nuclei (DAPI, blue). Scale bar: 50 µm. **(C)** Quantification of fusion index and area occupied by multinucleated osteoclasts (MNOCL, ≥3 nuclei) (n=4 independent experiments, 4 images/experiment, at least 1,000 nuclei/image, mean +/− SD). Quantification of nuclei number per multinucleated osteoclasts (one representative experiment, 150-250 cells/condition, from n=3). (**D-E)** Analysis of TNT formation. **(D)** Quantification of the percentage of cells forming thick (defined as positive for both F-actin and microtubules) and thin (defined as positive for F-actin only) TNTs on day 3 of differentiation after immunofluorescence analysis (see methods and panel E), n=4 independent experiments, n>250 cells/condition/experiments. Data are presented as mean ± SD, with each dot representing one independent experiment. Statistical analyses are performed on thick TNTs compared to controls only. **(E)** Representative microscopy images. Top panels: F-actin (phalloidin, gray, inverted). Bottom panels: F-actin (magenta), nuclei (DAPI, blue) and microtubules (MTs, α-tubulin, green). Scale bar: 20 μm.

Functional assessment of these mutant cells revealed distinct effects on osteoclast fusion and multinucleation (Figure 2B). Compared to control osteoclasts, deletion of Talin1 resulted in a marked reduction in the number of adherent cells, accompanied by a pronounced decrease in the fusion index, the surface area occupied by multinucleated cells, and the mean number of nuclei per cell. Collectively, these observations indicate a near-complete inhibition of osteoclast fusion in the absence of Talin1 (Figure 2C). Since Talin1 is a central regulator of integrin activation and binding to the actin cytoskeleton, its deletion closely replicated the phenotype of combined loss of β1, β2, and β3 integrins in osteoclasts during osteoclast fusion (Figure S3A-C). Consistent with previous findings, deletion of integrin β2 alone did not significantly alter fusion, while loss of integrin β1 tended to increase the fusion index and loss of β3 to decrease it, as previously described and in agreement with its role as a marker of osteoclast differentiation (Schmidt *et al*., 2011; Teitelbaum, 2011). These observations support the concept of partially redundant yet distinct roles of integrins in osteoclast formation, effects that can be reproduced by deletion of Kindlin3, similarly to Talin1 (Schmidt *et al*., 2011; Zou *et al*., 2013).

Loss of Lpd expression in osteoclast precursors leads to a highly significant impairment in cell fusion and reduced the formation of mature multinucleated osteoclasts. By contrast, RIAM deficiency produces a more moderate phenotype, characterised by the formation of smaller osteoclasts containing fewer nuclei, without a statistically significant reduction in the fusion index (Figure 2B-C). We further evaluated the formation of actin-positive TNTs, distinguishing thin TNTs lacking microtubules from thick TNTs containing microtubules, which participate in the fusion of osteoclast precursors (Dufrançais *et al*., 2025). Consistent with these observations, we found a strong correlation between the reduced number of thick TNTs and impaired fusion in Talin1- and Lpd-deficient cells, while RIAM deficiency did not significantly affect their abundance. In contrast, the number of thin TNTs remained relatively stable across all conditions. Together, these results identify Talin1 and Lpd as key regulators of osteoclast precursor fusion, likely through their role in the formation of functional thick TNTs, while RIAM appears to play a more limited or modulating role in this process.

### RIAM and Lpd contribute to osteoclast-mediated bone resorption

The capacity of osteoclasts to degrade bone is generally correlated with their degree of multinucleation and their overall cell size. Given the distinct roles of RIAM and Lpd in the fusion of osteoclast precursors, and their preferential localisation within the podosome belt when cells are cultured on glass, we next assessed their function under more physiological conditions by differentiating and culturing osteoclasts on bone substrates. Bone resorption was visualised by toluidine blue staining, revealing degradation pits and trenches generated by osteoclast activity (Figure 3A). Quantitative analysis of the resorbed surface area demonstrated a significant reduction in the bone-degrading capacity of RIAM- and Lpd-deficient osteoclasts compared to control cells (Figure 3B). However, this impairment was less pronounced than that observed after Talin1 deletion, which results in a near-complete inhibition of resorption activity, as previously reported (Zou *et al*., 2013). The critical role of Talin1 is all the more evident given that bone resorption is significantly reduced upon the simultaneous loss of β1, β2, and β3 integrins, unlike the deletion of each integrin subunit individually. Indeed, the loss of β1 or β2 alone does not significantly affect the resorption capacity of osteoclasts, while the deletion of β3 results in only a partial defect (Figure S3D-E). These findings are consistent with previous studies demonstrating the functional redundancy of integrins in podosomes and the formation of the sealing zone, as well as their collective contribution to bone degradation (Schmidt *et al*., 2011). Taken together, these results indicate that RIAM and Lpd both contribute to the bone resorption activity of osteoclasts, although their effects are less decisive than that of Talin1, thus highlighting their modulatory role in osteoclast function.

**Figure 3:**
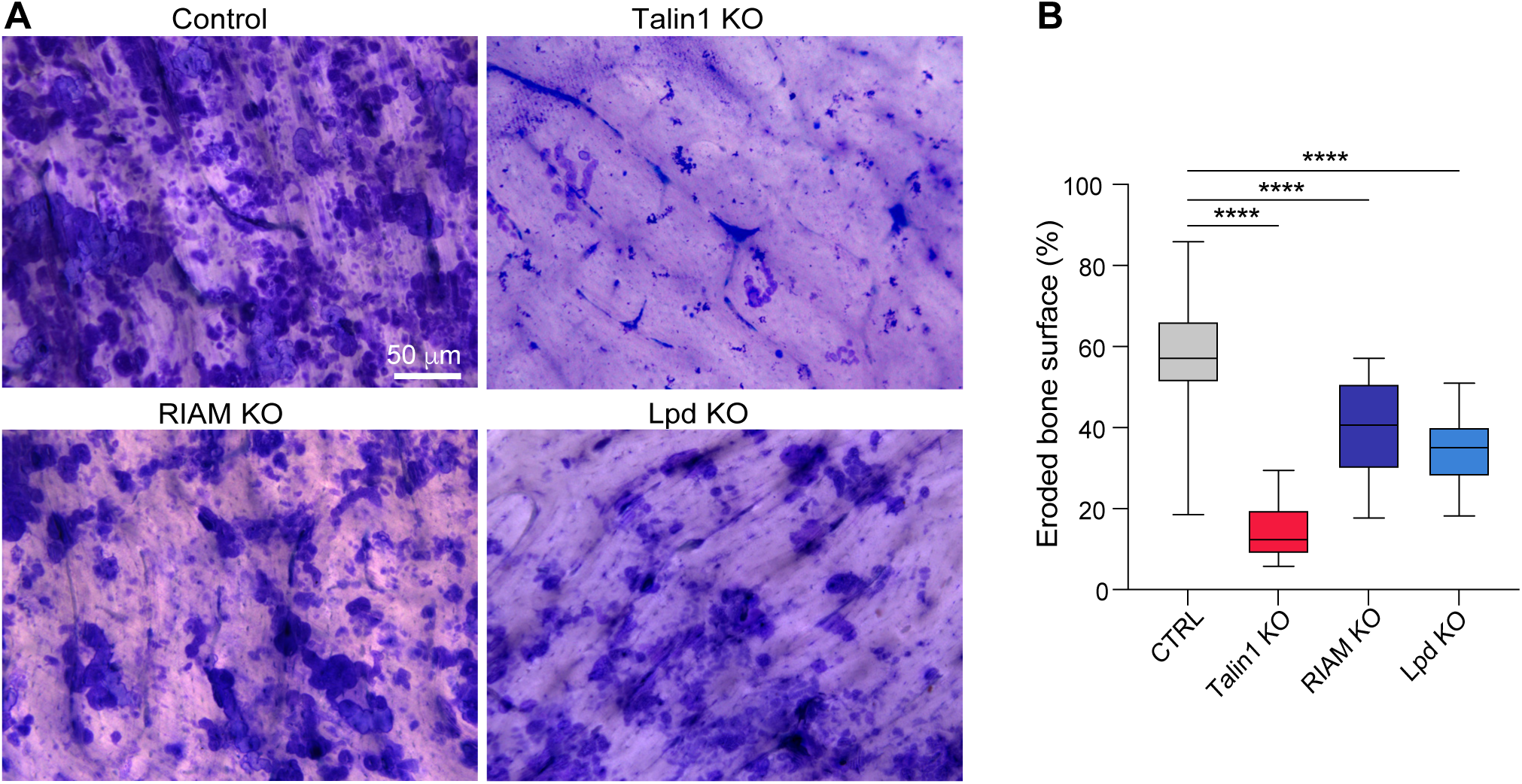
RIAM and Lpd deletion affects bone degradation in murine osteoclasts. (A-B) Effect of Talin1, RIAM or Lpd KO on bone degradation by osteoclasts cultured for 7 days on bone slices; after cell removal, the bones were stained with toluidine blue. **(A)** Representative images of bone degradation, eroded bone surfaces appear blue. Scale bar: 50 µm. **(B)** Quantification of eroded bone surface area (%) by semi-automatic quantification (n=3 independent experiments, 2 or 3 bone slices per condition, means ± SD are shown).

### Disruption of the Rap1–Talin1 interaction impairs osteoclast fusion and bone resorption

Deletion of Rap1 at late stages of osteoclast differentiation, under the control of the *CtsK* promoter, generates osteoclasts with a crenulated morphology (Zou *et al*., 2013). This phenotype is characteristic of cells lacking integrin-activated cytoskeletal organisers, particularly Talin1, and is associated with impaired bone resorption. In addition to signalling through its effectors RIAM and Lpd, Rap1 also binds directly to the F0 and F1 subdomains within the N-terminal head domain of Talin1. To evaluate the contribution of this direct interaction, we analysed osteoclast differentiation in cells carrying the R35E and R118E point mutations of Talin1 (Figure S4), hereafter referred to as Talin1 RBM (Rap1-binding mutant), which prevent binding to Rap1 (Gingras *et al*., 2019; Lagarrigue *et al*., 2020). RNA-seq analysis revealed no significant differences in gene expression between control osteoclasts and those expressing the Talin1 RBM mutant (not shown). Further proteomic analysis confirmed that the mutant Talin1 RBM protein is expressed at levels comparable to those of the wild-type protein, while wild-type specific are absent in Talin1 RBM-expressing cells, as expected (Figure S5A-B). In addition, parallel differentiation of the same progenitors into osteoclasts and macrophages, followed by global proteomic profiling, showed an enrichment in key osteoclast markers (Figure S5C). Overall, no significant changes were observed in the expression of proteins essential for osteoclast differentiation and function, including NFATc1, DC-STAMP, CtsK, and MMP9, nor in that of integrin subunits such as αvβ3, or in that of the proteins of interest Talin1, Rap1 (a and b isoforms), RIAM, and Lpd (Figure S5D-E). These findings validate the experimental model and indicate that mutations in the Talin1 RBM protein do not broadly affect osteoclast differentiation programmes neither integrin expression and regulation.

After differentiation, osteoclasts carrying the Talin1 RBM protein exhibit normal adhesion and spreading when cultured on vitronectin (Figure 4A-B). However, their ability to form TNTs is reduced by approximately 50% (Figure 4C), and their fusion capacity is significantly impaired (Figure 4D), leading to a marked decrease in the fusion index, the size of multinucleated osteoclasts, and the mean number of nuclei per cell on day 4 of differentiation (Figure 4E). Kinetic analysis shows that by day 5, mutant osteoclasts partially recover their fusion capacity (Figure 4D-E, Movie 1), reaching a fusion index comparable to that of control cells, while remaining smaller, as quantified in Figure 4E (day 5). This indicates that disruption of the Rap1–Talin1 binding delays, rather than completely blocks, osteoclastogenesis. Talin1 RBM-expressing osteoclasts also displayed a significant reduction in bone resorption. Toluidine blue staining revealed a marked decrease in resorbed surface area (Figure 5A-B), which was corroborated by a ∼50% reduction in β C-terminal telopeptide (CTx) levels—a marker of collagen degradation—in culture supernatants (Figure 5C). Furthermore, F-actin staining of osteoclasts detached and seeded on bone substrates for 3 days showed defective formation and organisation of adhesion structures. Cells expressing Talin1 RBM largely failed to establish sealing zones, and when these structures were present, they were smaller and disorganised than that observed in control cells (Figure 5D-E). Together, these results demonstrate that the direct interaction between Rap1 and Talin1 is required for efficient osteoclast fusion, adequate cytoskeletal organisation, and optimal bone resorption activity.

**Figure 4:**
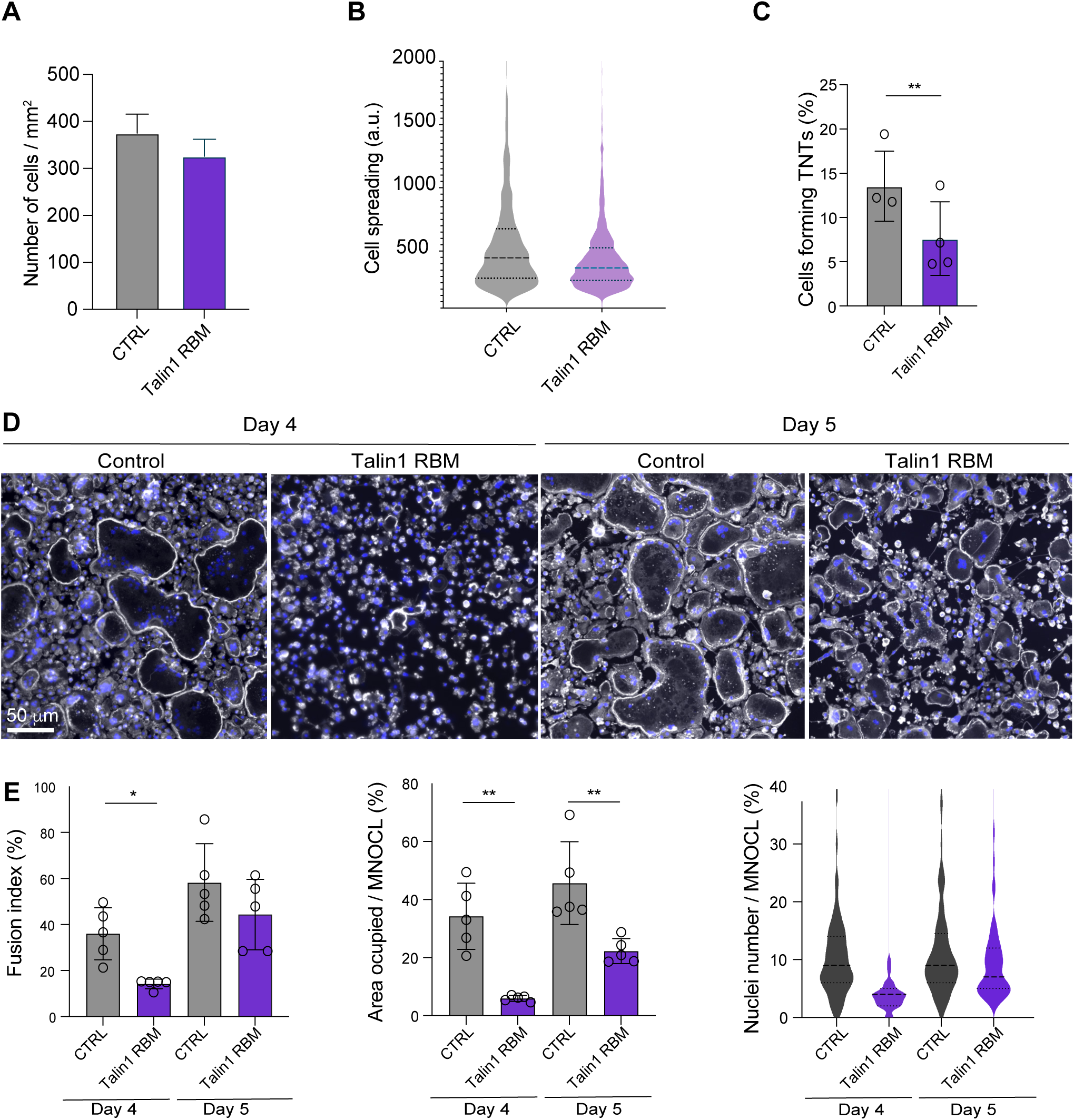
Consequences of disruption of the Talin1-Rap1 interaction (Talin1 RBM) in murine osteoclasts. (**A-B)** Analysis of osteoclast adhesion **(A)** and spreading **(B)** on vitronectin on day 3 of differentiation. **(C)** Analysis of TNT formation. Quantification of the percentage of cells forming thick TNTs on day 3 of differentiation after immunofluorescence analysis (see methods), n=4 independent experiments, n>250 cells/condition/experiments, mean ± SD are shown. **(D-E)** Analysis of cell fusion. Microscopy analysis of cell fusion in control and Talin1 RBM osteoclasts on days 4 and 5 of differentiation. **(D)** Representative microscopy images: F-actin (phalloidin, white) and nuclei (DAPI, blue). Scale bar: 50 µm. **(E)** Quantification of the fusion index and the surface area occupied by multinucleated osteoclasts (MNOCL, ≥3 nuclei) (n=5 independent experiments, 4 images/experiment, 1,000 nuclei/image, data are shown as means ± SD with each dot representing the mean of one independent experiment.). Quantification of the number of nuclei per multinucleated osteoclasts (one representative experiment, 150-250 cells/condition, from n=4).

**Figure 5:**
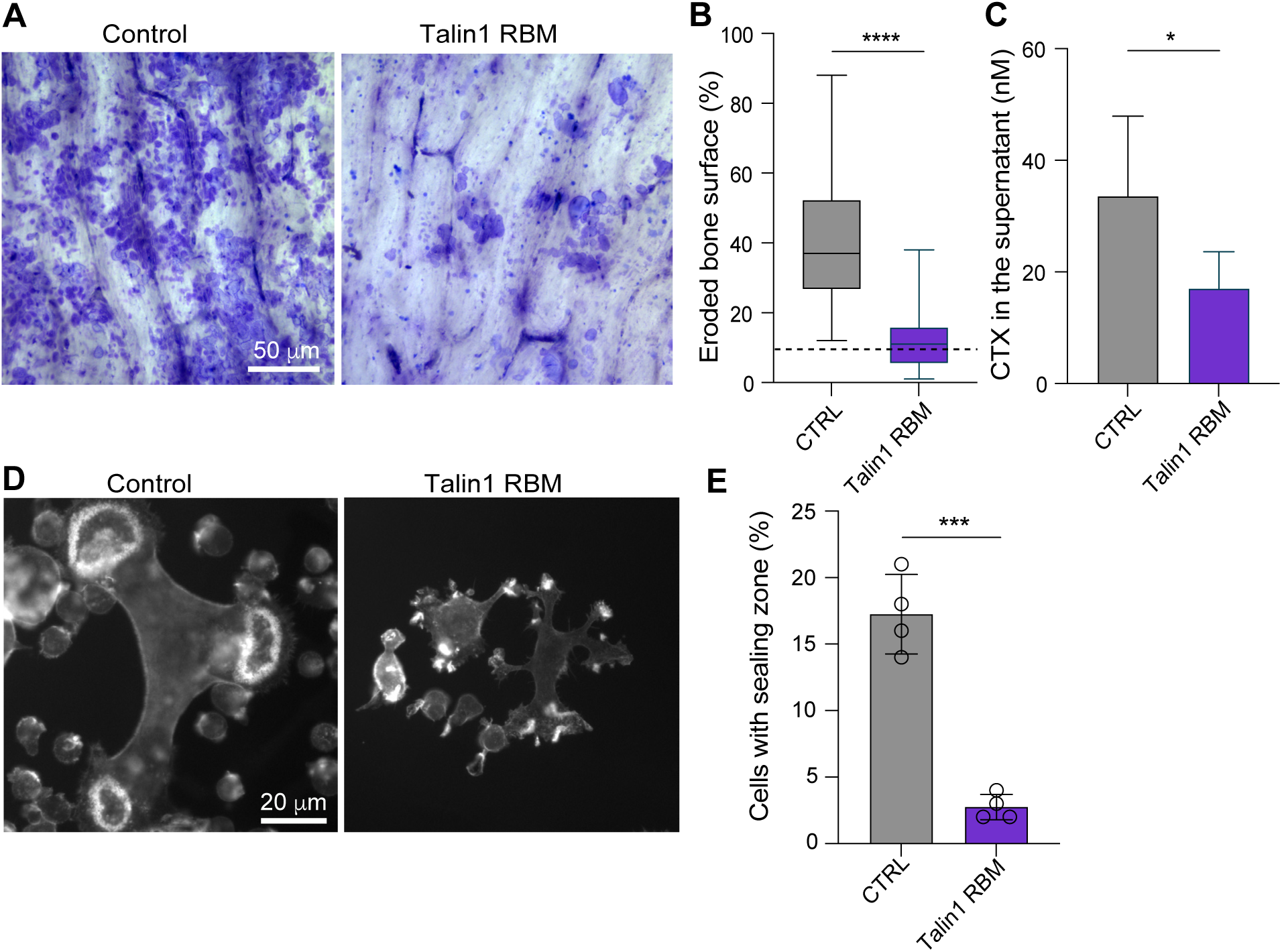
Disruption of the Talin1-Rap1 interaction (Talin1 RBM) affects bone degradation and sealing zone formation in murine osteoclasts. Effect of the disruption of Talin1-Rap1 interaction (Talin1 RBM) on bone degradation **(A-C)** and sealing zone formation **(D-E)**. **(A-B)** Osteoclasts were cultured for 7 days on bone slices, after supernatant recovery and cell removal, the bones were stained with toluidine blue. **(A)** Representative images of bone degradation, eroded bone surfaces appear blue. Scale bar: 50 µm. **(B)** Quantification of eroded bone surface area (%) by semi-automatic quantification (n=4 independent experiments, 2 or 3 bone slices/condition, means ± SD are shown). **(C)** Quantification of CTX (nM) by ELISA (n=4 independent experiments, means ± SD are shown). **(D-E)** Osteoclasts differentiated on day 3 were detached and cultured for 3 days on bone slices; the cells were then fixed. **(D)** Microscopy images representing sealing zones visualized by F-actin staining (phalloidin, white). Scale bar: 20 µm. **(E)** Quantification of the number of cells forming sealing zones, n=4 independent experiments, n>250 cells/condition/experiments, means ± SD are shown.

### Direct disruption of the Rap1–Talin1 interaction, but not the loss of RIAM or Lpd, alters bone homeostasis

To investigate the physiological relevance of disrupting the functional interactions linking Rap1 to Talin1 in osteoclasts and bone biology, we generated mouse models carrying conditional inactivation of the genes encoding RIAM (Klapproth *et al*., 2015) and Lpd (Law *et al*., 2013), using the Cre recombinase driven by the *LysM* promoter (Clausen *et al*., 1999). Since mutations in the Rap1-binding motifs (RBM) of Talin1 are lethal at the embryonic stage in the homozygous state (Lagarrigue *et al*., 2020), we combined the Talin1 RBM mutant allele with a floxed Talin1 allele. This approach restricts the expression of the mutant Talin1 protein to Cre-expressing myeloid lineage cells, including osteoclasts. Mice from the same litter, expressing the wild-type Talin1 allele, served as controls.

In all models, a fluorescent reporter gene eYFP, positioned downstream of a stop codon flanked by loxP sites, allowed for the identification of cells expressing Cre recombinase (and thus genetically modified). Immunohistochemical analysis of serial sections of femur from control mice showed that more than 85% of Cathepsin K-positive osteoclasts lining the bone surface express eYFP, as do other resident bone marrow cells, thereby validating the model (Figure 6A and Movie 2). The osteoclastic identity of these eYFP-positive cells was confirmed by their multinucleation, elongated morphology, and localisation on the bone surface.

**Figure 6:**
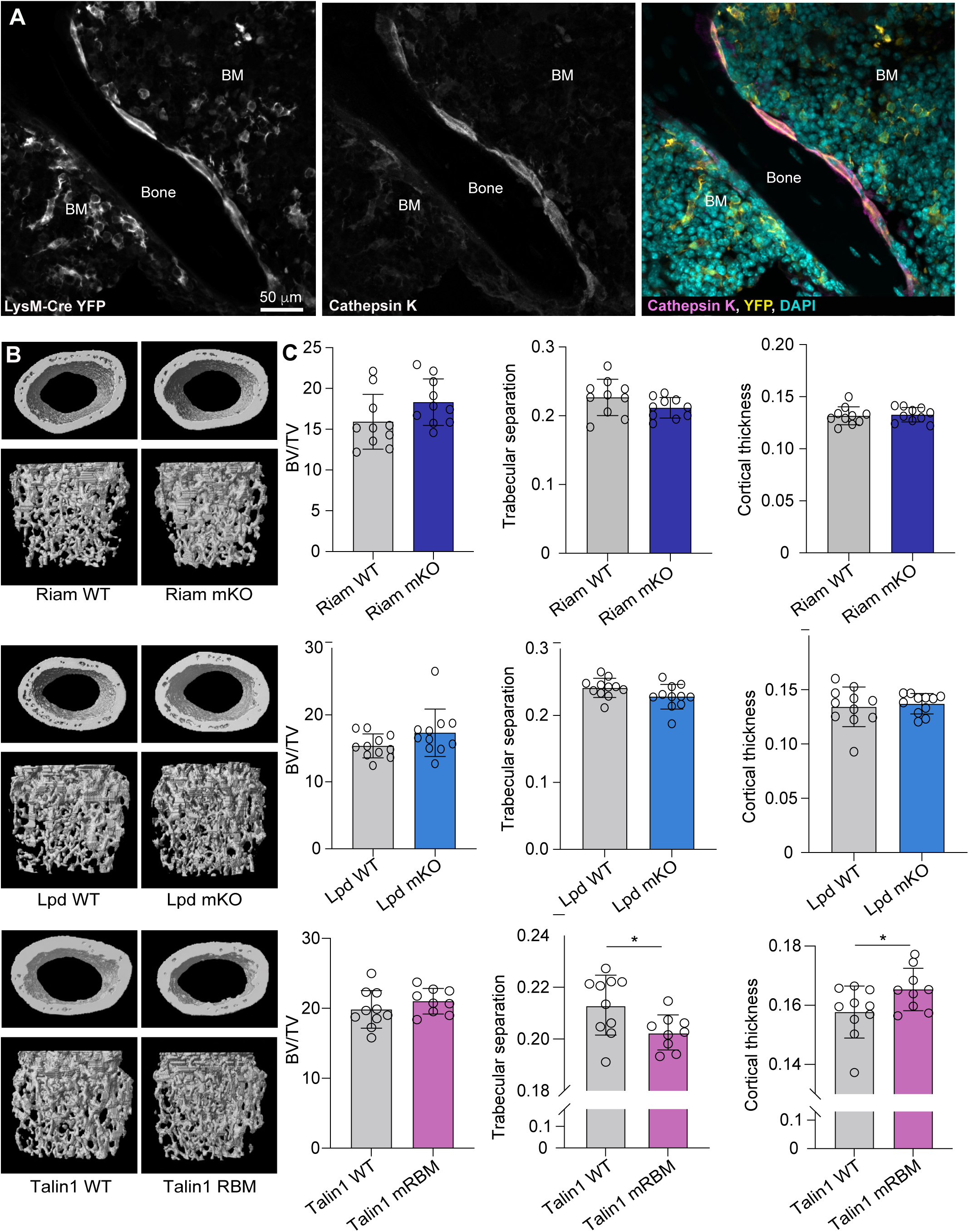
Disruption of the Talin1-Rap1 interaction results in moderate bone defects in vivo. **(A)** Representative confocal image of the histological analysis of wild-type (*LysM^Cre/+^; R26R^eYFP/eYFP^*) mouse femora. nuclei (DAPI, cyan), and eYFP (yellow) or Cathepsin K (pink). Scale bar: 50 µm. BM: Bone marrow. See also Movie 2. **(B)** Representative X-ray microtomography images of cortical (upper panels) and trabecular (lower panels) sections of distal femora from wild-type mice and Lpd mKO, RIAM mKO or Talin1 mRBM mice. Scale bar: 500 µm. **(C)** The histograms represent the mean ± SD of trabecular bone volume per total volume fraction (left), trabecular separation (center) and cortical thickness (right). Each group of animals consisted of 10 mice.

We next assessed the skeletal phenotype of these mice. No differences in body size, weight, or overall skeletal architecture were observed between genotypes up to 16 weeks of age (data not shown). X-ray microtomography analysis of the distal femurs of 12-week-old mice revealed that conditional knockout of RIAM or Lpd did not significantly alter trabecular or cortical bone parameters (Figure 6B-C and S6). In contrast, Talin1 RBM mutant mice exhibited a slight but significant decrease in trabecular separation compared to controls, accompanied by a slight increase in cortical bone thickness, without any change in trabecular bone volume (Figure 6B-C). Analysis of tartrate-resistant acid phosphatase (TRAP) staining revealed a significant reduction in osteoclast activity in Talin1 RBM mutant mice, at a level comparable to that observed after conditional deletion of Talin1 (Figure 7A). Immunohistochemical analysis of Cathepsin K-positive osteoclasts in femoral sections further demonstrated that the Talin1 RBM mutation, similarly to Talin1 loss, leads to a reduction in the number and /or activity of osteoclasts, while their abundance remains unchanged in mice lacking RIAM or Lpd (Figure 7). Taken together, these results show that disruption of the direct Rap1–Talin1 interaction specifically alters the number and activity of osteoclasts in vivo, leading to measurable changes in bone microarchitecture, whereas loss of RIAM or Lpd alone has no detectable impact under these conditions.

**Figure 7:**
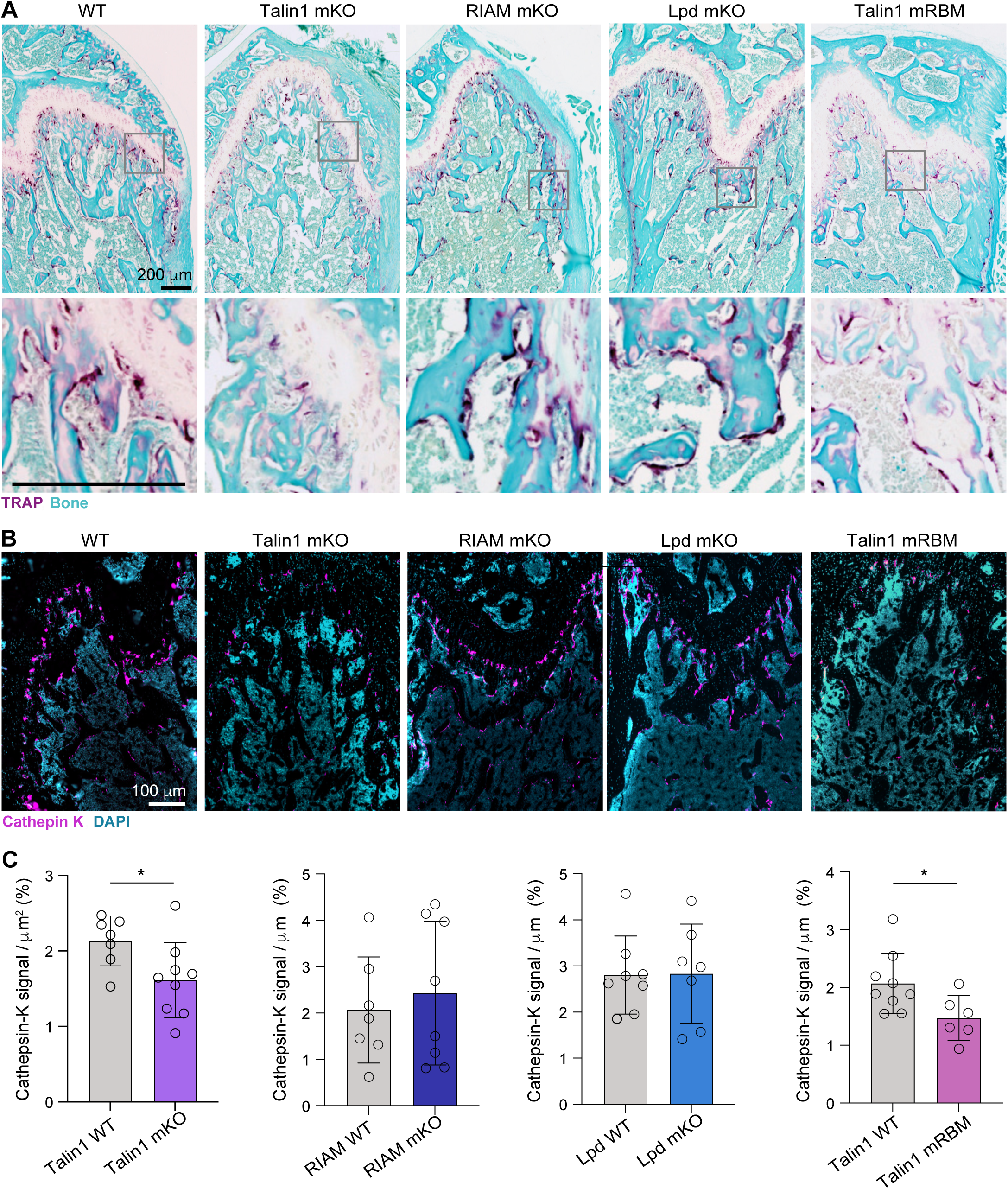
Disruption of the Talin1-Rap1 interaction reduces osteoclast activity in vivo. **(A-C)** Histological analysis of osteoclasts by TRAP **(A)** and Cathepsin K **(B-C)** staining in femora from wild-type mice and Lpd mKO, RIAM mKO, or Talin1 mRBM mice. **(A)** Representative microscopy image of the histological analysis TRAP-stained femora counterstained with fast green. Scale bar: 200 µm. **(B)** Representative image of the fluorescence microscopy histological analysis of femurs stained with Cathepsin K (magenta) and DAPI (nuclei, blue). Scale bar: 100 µm. **(C)** The histogram represents the mean ± SD of Cathepsin K signal area per unit bone area for each condition. n=3-4 mice, 2-3 images/bone.

## Materials and methods

### Mice

All animal experiments were conducted in accordance with national and European regulations and institutional guidelines for animal studies. Mice were housed in the animal facilities of IPBS, part of the ANEXPLO/Genotoul life sciences platform in Toulouse, under conditions free of specific pathogens. The Cre driver strains under the control of the LysM (Clausen *et al*., 1999)or CMV (Schwenk, Baron and Rajewsky, 1995) promoters and the R26R^eYFP/eYFP^ strain (Srinivas *et al*., 2001) come from the Jackson laboratory. Mice carrying floxed alleles of the *Tln1* (encoding Talin 1) (Petrich *et al*., 2007), *Apbb1ip* (encoding RIAM) (Klapproth *et al*., 2015; Su *et al*., 2015), or *Raph1* (encoding Lpd) (Law *et al*., 2013) genes have been described previously. The *Tln1* allele carrying the R35E and R118E mutations, which block the direct binding of the Talin1 protein to Rap1, hereafter referred to as RBM for Rap1 binding defective mutant, has been described previously (Lagarrigue *et al*., 2020). Cas9-expressing mice have also been described (Tzelepis *et al*., 2016). Each strain was backcrossed more than five times on a C57BL/6J genetic background before any crossing.

Conditionally knockout mouse strains were generated by crossing floxed mice with the *LysM^Cre/+^* strain and *R26R^eYFP/eYFP^* mice that possess a STOP sequence flanked by loxP followed by the eYFP gene inserted in the *Gt(ROSA)26Sor* locus so that eYFP serves as a marker of Cre recombinase activity and enables the labelling of target cells. Thus, same-sex *Tln1^flox/flox^; R26R^eYFP/eYFP^; LysM^+/+^* mice served as controls for *Tln1^flox/flox^; R26R^eYFP/eYFP^; LysM^Cre/+^* mice (designated Talin1 mKO, corresponding to specific myeloid cell inactivation). The same approach was applied to the *Apbb1ip* and *Raph1* strains, hereinafter referred to as RIAM mKO and Lpd mKO respectively. In the specific case of mice carrying the R35E and R118E mutations of the *Tln1* gene, we crossed *Tln1^RBM/+^; R26R^eYFP/eYFP^* mice with the *Tln1^flox/flox^; R26R^eYFP/eYFP^; LysM^Cre/+^* strain to obtain a specific myeloid deletion of the Tln1-Flox allele in *Tln1^flox/+^; R26R^eYFP/eYFP^; LysM^Cre/+^*(control) and *Tln1^flox/RBM^; R26R^eYFP/eYFP^; LysM^Cre/+^*(Talin1 mRBM) mice from the same litter, for the experiments.

To induce global suppression of the *Apbb1ip* or *Raph1* gene, floxed mice were crossed with the CMV-driven Cre strain to induce germline suppression, then backcrossed with wild-type C57BL6/J mice to eliminate the Cre gene. Given the postnatal lethality of *Raph1^−/−^*newborns, as previously described (Law *et al*., 2013), we maintained this strain in a mixed C57BL/6J and FVB/N genetic background. None of the mutant mouse strains exhibited major developmental defects or signs of distress.

### HoxB8-derived murine osteoclasts

Myeloid progenitors were isolated from mouse bone marrow and immortalized by transduction using a retrovirus conditionally expressing the HoxB8 homeotic gene. Specifically, bone marrow cells were collected and hematopoietic progenitor cells were purified by centrifugation on a Ficoll-Paque Plus (Cytiva). These cells were then pre-stimulated for two days in complete medium [RPMI 1640 GlutaMAX containing 10% foetal bovine serum (FBS) and 1% v/v antibiotic-antifungal (Gibco), supplemented with murine IL-3, murine IL-6, and murine SCF, each at a concentration of 10 ng/mL (Miltenyi-Biotec). Next, 2 × 10⁵ cells/mL were seeded into a six-well culture plate containing myeloid medium [complete medium, supplemented with 20 ng/mL murine GM-CSF (Miltenyi-Biotec) and 0.5 μM β-estradiol (Sigma-Aldrich) and transduced with an MSCV retrovirus encoding the oestrogen-binding domain of the oestrogen receptor (ER) fused to the N-terminus of murine HoxB8. Retroviral particle production occurred over three days after transfection of HEK293 cells with the pMSCV and pCL-Eco plasmids (1:1 ratio), performed using the JetOptimus transfection reagent (Polyplus), according to the manufacturer’s instructions. The supernatant from virus-producing HEK293 cells was used and diluted 1:5 for transduction with Lentiblast Premium (OZ Biosciences). One to two weeks after transduction, only immortalized ER-HoxB8 progenitors continued to proliferate. These progenitors were then sub-cultured every two days in myeloid medium maintained at a concentration of 5 × 10^4^ to 1 × 10⁶ cells/mL.

To generate cell lines with knockouts for the *Tln1*, *Apbb1ip*, or *Raph1* genes, we immortalized myeloid progenitors from floxed but Cre-deficient mice. After immortalization, we transduced them with a lentivirus containing the cassette expressing Cre-IRES-mCherry, or mCherry alone as a control, downstream of the murine PGK promoter cloned between the Cla1 and Csi1 restriction sites of the pLVX-Puro vector (Takara). Lentiviral particles were produced by transfection of HEK293 cells with the plasmids pMDL (Addgene #12251), pREV (Addgene #12253), and pVSV-G (Addgene #12259), as well as with cloned pLVX plasmids, in a 2:1:1:4 ratio, using JetOptimus reagent (Polyplus) according to the manufacturer’s instructions. The supernatant from the virus-producing HEK293 cells was collected 3 days post-transfection and diluted 1:5 to transduce 2 × 10⁵ cells/mL with Lentiblast Premium (OZ Biosciences). Cells doubly positive for mCherry and eYFP (whose expression is induced by Cre activity) were sorted by flow cytometry (BD FACSAria™ Fusion) to obtain knockout cells, while their wild-type counterparts, used as controls, were sorted solely for mCherry expression (without Cre expression). In the specific case of the Talin1 RBM mutation, both control cells (*Tln1^flox/+^; R26R^eYFP/eYFP^*) and mutant cells (*Tln1^flox/RBM^; R26R^eYFP/eYFP^*) were transduced by the Cre-mCherry construct and sorted on the basis of double-positive cells for mCherry and eYFP.

For the differentiation of osteoclasts, HoxB8 progenitors were collected, washed twice, and seeded onto glass coverslips (Paul Marienfeld GmbH) in complete medium supplemented with 25 ng/mL of murine M-CSF (Miltenyi-Biotec) and 100 ng/mL of murine RANKL (R&D Systems), at a rate of 1 × 10⁵ cells per well of a 24-well plate and 1 × 10⁵ cells per well of a 6-well plate. The cells were incubated at 37 °C in a 5% CO₂ incubator. The medium and cytokines were renewed every 3 days, and mature osteoclasts were obtained between the 5th and 7th day.

### CRISPR-Cas9-mediated gene editing in HoxB8 progenitors

Gene inactivation was performed using CRISPR-Cas9 technology. Myeloid progenitors from a Cas9-expressing mouse were immortalized by retroviral transduction of the ER-HoxB8 gene. Once established, the cells were transduced with lentiviruses encoding sgRNAs. The sgRNA sequences used were *Itgb1* (5’-gttggtcagcaacgcatatc-3’), *Itgb2* (5’-agactcggtgatctcgttga-3’) and *Itgb3* (5’-atgtacatgtacggcgatac-3’). Luciferase was targeted as a control (5′-ggcgcggtcggtaaagttgt-3′). For single-gene knockout, the sgRNAs were inserted into the pLenti-sgRNA vector (Addgene #71409) between the two BsmBI restriction sites. For the triple inactivation of the *Itgb1*, *Itgb2*, and *Itgb3* genes, encoding β1, β2, and β3 integrins, respectively, the three sgRNAs were cloned by Golden Gate assembly into the pLV hUbC-Cas9-T2A-GFP plasmid (Addgene #53190), in which the GFP gene was replaced with a puromycin resistance gene. Lentiviral particles carrying sgRNA were produced using the same procedure as previously described, as was lentiviral transduction of cells. The following day, the transduced cells were selected with 10 µg/mL of puromycin (Invivogen) for 2 days, and gene inactivation was confirmed by flow cytometry.

### Human monocyte-derived osteoclasts

Human peripheral blood mononuclear cells were isolated from leukocyte concentrates of blood from healthy donors (French National Blood Institute, contract 28 21/PLER/TOU/IPBS01/20130042). Written informed consent was obtained from all participants. The cells were centrifuged on a Ficoll-Paque Plus (Cytiva), resuspended in cold PBS supplemented with 2 mM EDTA and 0.5% heat-inactivated foetal bovine serum (FBS) at pH 7.4, and the monocytes were sorted using magnetic microbeads coupled to anti-CD14 antibodies (Miltenyi-Biotec). For differentiation into human osteoclasts, monocytes were seeded onto glass coverslips (Paul Marienfeld GmbH) in 24-well plates at a density of 5 × 10⁵ cells per well in RPMI GlutaMAX medium supplemented with 10% FBS, human M-CSF (50 ng/mL; Miltenyi-Biotec), and human RANKL (30 ng/mL; Miltenyi-Biotec). The cells were incubated at 37 °C in a 5% CO₂ incubator. The medium was renewed every 3 days with a medium containing M-CSF (25 ng/mL) and RANKL (100 ng/mL). Mature osteoclasts were obtained between days 7 and 10.

### Immunofluorescence

Cells were fixed with PFA (3.7%, Sigma-Aldrich), 30 mM sucrose in PBS (Gibco), permeabilized with 0.3% Triton X-100 (Sigma-Aldrich) for 10 min, and then blocked with BSA (1% in PBS) for 30 min. After incubation with the primary antibodies for 1 h, the cells were washed with PBS and then incubated with the corresponding fluorescent secondary antibodies (2 μg/mL, Jackson ImmunoResearch), fluorescently labelled phalloidin (Invitrogen), and DAPI (500 ng/mL, Sigma-Aldrich) for 30 min. After several washes with PBS, the coverslips were mounted on a glass slide using fluorescence mounting medium (Dako) and stored at 4°C. The primary antibodies used were: RIAM (1/100; AF7279; R&D systems), Lpd (1/100; 90859; Novus Biologicals), Talin1 (1/500; SAB4200694; Sigma-Aldrich), α-Tubulin (1/200, 047M4760V; Sigma-Aldrich). The fluorophore-conjugated secondary antibodies used were: donkey anti-rabbit IgG (1/500; 300-545-245; Jackson ImmunoResearch), donkey anti-goat IgG 1/500; 713-605-147; Jackson ImmunoResearch), donkey anti-mouse IgG (1/500; 715-575-151; Jackson ImmunoResearch). Phalloidin (1/400, Molecular Probes, Alexa 555 or 488) and DAPI (D9542; Sigma-Aldrich) were used to stain F-actin and nuclei, respectively.

To analyse osteoclast fusion, images were acquired with a Leica DMi8 inverted optical microscope equipped with a Plan Apochromat 20×/0.8 objective and a K5 camera, controlled by LasX software (Leica Microsystems). The fusion index is defined as the number of nuclei present in a multinucleated giant cell (> 3 nuclei) relative to the total number of nuclei per field (Vérollet *et al*., 2015; Raynaud-Messina *et al*., 2018) The osteoclast fusion index, the number of nuclei per multinucleated cell, and the surface area occupied by these cells were quantified using a semi-automated method with a specifically developed ImageJ macro. For each condition, 4 images and at least 1,000 nuclei per image were analysed.

Thin and thick tunnelling nanotubes (TNTs) were identified by phalloidin and α-tubulin staining and counted on at least 200 cells per condition, as performed previously (Dufrançais *et al*., 2025). We quantified TNTs as F-actin-positive structures that connect at least two cells and that do not adhere to the glass slides. Thick TNTs were classified compared to thin counterparts by their diameter (≥2 nm) and the presence of microtubules. The number of cells forming sealing zones were quantified after phalloidin staining of mature osteoclasts cultured on bone slices. Most of the images were acquired using a Zeiss Axio Imager M2 microscope equipped with a Plan Apochromat 20×/0.8 or 40×/0.95 objective (Zeiss) and an ORCA-flash 4.0 LT camera (Hamamatsu). Confocal images were acquired using a Zeiss LSM880 inverted laser scanning confocal microscope equipped with a Plan-Apochromat 63×/1.4 NA oil immersion objective. Z-stacks were collected at optimal intervals and mage acquisition and processing were performed using ZEN software (Zeiss), and then using ImageJ and Photoshop software.

To visualize sealing zone formation, osteoclasts were differentiated on glass coverslips for three days, detached using Accutase, and subsequently reseeded onto bone slices for an additional three days. Cells were then fixed and stained with phalloidin to visualize F-actin organization.

### Bone resorption assay

To assess bone resorption activity, HoxB8 precursor cells were seeded onto 6 mm diameter and 0.2 mm thick bovine cortical bone slices (IDS Nordic Biosciences) in 96-well plates (5 × 10⁴ cells/well). Osteoclasts were differentiated in complete medium, supplemented with 50 ng/mL of M-CSF (Miltenyi-Biotec) and 30 ng/mL of RANKL (R&D Systems), for 10 days. After complete cell removal by several water washes, bone slices were stained with toluidine blue (Sigma-Aldrich) to detect resorption lacunae using a Leica DMi8 microscope equipped with a Fluotar 10×/0.32 objective and a K3C camera, controlled by LasX software (Leica Microsystems). The bone degradation surface area was quantified with a home-made semi-automatic macro using ImageJ software, as performed previously (Dufrançais *et al*., 2025). A minimum of 2 to 3 bone slices per condition were analysed and 3-6 images per bone.

### Flow cytometry

Cells were washed in PBS and collected by centrifugation at 500 × g for 5 min, then stained with the LIVE/DEAD kit (Thermo Fisher Scientific) according to the manufacturer’s instructions. Cells were counted and seeded in a 96-well plate at a density of 3 × 10⁵ cells/well. Surface staining was performed for 30 min at 4 °C in 100 μL of staining buffer (PBS containing 3% BSA) with antibodies conjugated to an appropriate fluorochrome, and then the cells were washed twice. For intracellular staining, cells were fixed using the Foxp3/transcription factor staining buffer kit (Thermo Fisher Scientific), according to the manufacturer’s instructions, before being stained with the appropriate primary and secondary antibodies for 1 h at 4 °C. They were then washed three times between and after the staining steps. Data were acquired using a BD Fortessa X20 flow cytometer (BD Biosciences) driven by BD FACS Diva software and analyzed using FlowJo (Tree Star). The antibodies used were: RIAM (1/100; AF7279; R&D systems), Lpd (1/100; 90859; Novus Biologicals), Talin1 (1/500; SAB4200694; Sigma-Aldrich Integrin β1 (1/200; 102207, Biolegend), Integrin β2 (1/200; 101414, Biolegend), Integrin β3 (1/200; 104318, Biolegend) and their appropriate isotype controls. The fluorophore-conjugated secondary antibodies used were: donkey anti-rabbit IgG (1/500; 300-545-245; Jackson ImmunoResearch), donkey anti-goat IgG 1/500; 713-605-147; Jackson ImmunoResearch), donkey anti-mouse IgG (1/500; 715-575-151; Jackson ImmunoResearch).

### Histological analysis

For immunohistofluorescence of frozen bone sections (10 μm, Leica CM1950), femur bones from 12-week-old mice were fixed in 4% PFA (Electron Microscopy Science) overnight at 4°C, decalcified for 10 days with 10% EDTA (Sigma-Aldrich) renewed daily, then incubated in 30% sucrose solution (Euromedex) overnight at 4°C before being embedded in OCT (CellPath). After saturation and permeabilization/blockage for 1 hour at room temperature with 10% donkey serum (Biowest S2170-100), and 0.2% Triton X-100 (Sigma-Aldrich), the cryosections were incubation overnight at 4°C with anti-GFP antibodies (1:200; 600-101-215; Rockland), and with a donkey anti-goat secondary antibody conjugated to Alexa Fluor 555 (1:400; No. A-11056; Invitrogen) at room temperature for 2 hours. In a second step, the cryosections were saturated with Mouse-On-Mouse (MOM) Blocking (vector Laboratories MKB-2213-1) for 1 hour, incubated with anti-cathepsin K antibodies (1:100; ab19027; Abcam) in MOM Protein Concentrate stock overnight at 4°C, and then with goat anti-mouse IgG2b secondary antibody conjugated to Alexa Fluor 647 (1:400; No. A21242; Invitrogen) in MOM Protein Concentrate stock at room temperature for 2 hours. Nuclei were stained with DAPI (Sigma-Aldrich). Confocal images were acquired using a Zeiss LSM880 inverted laser scanning confocal microscope equipped with a Plan-Apochromat 63×/1.4 NA oil immersion objective. Z-stacks were collected at optimal intervals and mage acquisition and processing were performed as described above.

For the quantification of the cathepsin K staining, 5µm paraffin sections were used after fixation, decalcification and rehydration, and only the second step of the protocol described above. Images were acquired using a Zeiss Axio Imager M2 microscope equipped with a Plan Apochromat X20/0.8 objective (Zeiss) and an ORCA-flash 4.0 LT camera (Hamamatsu), and then processed with Zeiss Zen software. The percentage of positive cathepsin K staining relative to surface area was quantified using QuPath software. Cathepsin K staining was quantified on at least two to three selected bone sections from three to four mice of each genotype.

For TRAP staining, femurs of mice were fixed, decalcified and rehydrated as previously described. Serial 5 μm longitudinal paraffin sections (5 μm) were stained for 30 minutes at 37°C in TRAP staining solution (TRAP basic incubation medium, Napthol AS-MX, Fast Red Violet LB salt) and fast green (F7252; Sigma-Aldrich) for 30 s. Images were acquired using a Zeiss Axio Imager M2 microscope equipped with a Plan Apochromat X20/0.8 objective (Zeiss) and an AxioCam 503 color camera (Zeiss), and then processed with Zeiss Zen software.

### Bone histomorphometric analysis

The leg bones of 12-week-old female mice were fixed overnight at 4°C in PBS supplemented with 4% PFA (Electron Microscopy Science), then washed and stored in 70% ethanol. Bone microarchitecture analysis by high-resolution X-ray microtomography (μCT) was performed using a high-resolution in vivo X-ray microtomography system (Quantum FX, Rigaku, Life Sciences, Revvity) hosted on the life Imaging Facility “Plateforme des Imageries du Vivant”, (UMR-S 1333 Santé Orale, Université Paris Cité and Sorbonne Paris Nord, Inserm, Santé Orale, F-92120 Montrouge, France). Standard acquisition parameters were applied (voltage 90 kV, current 160 µA and Aluminium 0.1mm Copper 0.06mm Acrylic 2mm filter). Three-dimensional acquisitions were performed with a 5 mm field of view, centred on the proximal metaphysis and mid-diaphysis of the femur, with an acquisition time of 240 s and a voxel size of 10 μm³ in the regions of interest (ROI). Image acquisition and microarchitectural analysis were performed according to published recommendations (Bouxsein *et al*., 2010).

Trabecular bone was analysed at the proximal metaphysis of the femur for each sample, while cortical bone was analysed at the mid-diaphysis. For image analysis, the datasets were first reoriented using DataViewer software (SkyScan, version 1.6.0.0; Kontich, Belgium) to standardize alignment and define the regions of interest (ROI). The ROIs were adjusted to the size of the femur and fixed at a height of 200 slices. Bone microarchitecture (BMA) quantification was performed using CT-Analyzer software (SkyScan, version 1.20.8.0). After image segmentation, three-dimensional microarchitectural parameters were calculated for the trabecular and cortical compartments: bone volume fraction (BV/TV; %), number of trabeculae (Tb.N; mm⁻¹), trabecular separation (Tb.Sp; mm), trabecular thickness (Tb.Th; mm), cortical surface area fraction (Ct.Ar/Tt.Ar; %), cortical thickness (Ct.Th; mm), and cortical porosity (Ct.Po; %). Three-dimensional volumetric renderings were generated for the trabecular and cortical bone using CTVol software (SkyScan, version 2.2.3.0).

### Global proteomic analysis by mass spectrometry

Osteoclasts differentiated in a 24-well plate were washed twice with PBS and lysed in 150 µL of lysis buffer containing 50 mM ammonium bicarbonate and 4% sodium deoxycholate (SDC), then boiled for 5 min at 95 °C. Each condition was performed in technical quadruplicates in the same experiment. DNA was degraded by the addition of nuclease (Pierce). Disulfide bridges and cysteine residues were reduced and alkylated by incubation with 10 mM Tris (2-carboxyethyl) phosphine hydrochloride (NeoBiotech) and 40 mM 2-chloroacetamide (Sigma-Aldrich) for 5 min at pH 8.5 and 95 °C. The samples were then subjected to tryptic digestion according to the SP3 method (Hughes *et al*., 2019) on Sera-Mag™ magnetic beads modified with carboxylate groups (Cytiva). The magnetic beads were added to the cell lysate in a 10:1 ratio, and binding was induced by the addition of ethanol to a final concentration of 50%. The supernatant was discarded, and the beads were washed four times by resuspension in 80% ethanol. The proteins were then eluted and digested by resuspension of the beads in 50 mM ammonium bicarbonate containing 5% trypsin (Promega) at 37°C overnight.

Approximately 50 ng of the obtained peptides were loaded onto Evotips (Evosep) and analysed by nanoLC-MS/MS using an Evosep One nanochromatography system coupled to a TIMS-TOF SCP mass spectrometer (Bruker). Peptide separation was performed on an Aurora Elite C18 column (15 cm × 75 µm × 1.7 µm, IonOpticks) with a Whisper Zoom 40SPD gradient (solvent A: 0.1% formic acid in water; solvent B: 0.1% formic acid in acetonitrile), at a flow rate of 150 nL/min. Mass spectra were acquired in DIA-PASEF mode over the precursor mass range [400–1000] m/z and for an ionic mobility of 1/K₀ [0.64–1.37]. The acquisition protocol consisted of eight consecutive TIMS ramps with an accumulation time of 100 ms, each with three MS/MS acquisition windows of 25 Th. The total cycle time was 0.96 s. The collision energy was increased linearly with respect to ionic mobility, from 59 eV at 1/K₀ = 1.6 V·s·cm⁻² to 20 eV at 1/K₀ = 0.6 V·s·cm⁻².

Raw data were analyzed and quantified using Spectronaut 20.2 in DirectDIA mode, against mouse reference sequences from the UniProt database (mouse reference proteome, 55,307 UniProt sequences, downloaded January 10, 2026). Carbamidimethylation (C) was defined as a fixed modification, and acetylation (N-terminal end of the protein) and oxidation (M) as variable modifications in the Pulsar search. Trypsin specificity was fixed to cleavage after K or R, peptide length was between 7 and 30 amino acids, and missed cleavages were tolerated. Validation was performed with a false positive rate (FDR) of 1% for both the precursor and the protein.

The results of differential protein expression were analysed using the R software. Volcano plots were generated to visualize proteins exhibiting differential abundance. For certain proteins of interest, normalized expression values were scaled (Z-score transformation) and visualized as heatmaps. An overrepresentation analysis (ORA) was performed to identify enriched pathways among the differentially expressed proteins. Protein–protein interaction networks were explored using the STRING software.

### Statistical analyses

All statistical analyses were performed using GraphPad Prism 9 software (GraphPad Software Inc.). Paired and unpaired two-tailed t-tests were applied to data following a normal distribution (determined by the Kolmogorov-Smirnov test), while two-tailed (unpaired) or paired (Wilcoxon test) t-tests were used in other cases. When multiple comparisons were performed, the statistical analyses used are detailed in the legend of the corresponding figure. We used the Friedman test followed by Dunn’s test, a one-way ANOVA followed by Tukey’s test, or a Kruskal-Wallis test followed by Dunn’s test for multiple comparisons, depending on the distribution of the data (determined by the Kolmogorov-Smirnov test). The threshold for statistical significance was set at p < 0.05 (*, p≤0.05; **, p≤0.01; *** p≤0.001; **** p≤0.0001).

## Discussion

The roles of Rap1 and Talin1 in osteoclastogenesis and osteoclast-mediated bone resorption have been previously characterised (Zou *et al*., 2013). In the present study, we examined Rap1 effectors known to regulate Talin1-dependent integrin activation—a process tightly coupled to the actin cytoskeleton—whose functions in osteoclasts remain poorly understood. We demonstrate that RIAM and Lpd are expressed in osteoclasts, localise to actin-rich structures such as TNTs and podosomes, and contribute to the structural organisation of mature cells. Functional analyses revealed distinct roles for these proteins. Lpd deficiency markedly impaired the fusion of osteoclast precursors, partly due to diminished TNT formation at early stages, resulting in reduced multinucleation, and a significant decrease in bone resorption. In contrast, RIAM deficiency had minimal impact on cell fusion but reduced the resorption capacity of mature osteoclasts, indicating a role at later stages. Disruption of the direct Rap1– Talin1 interaction produced a broader phenotype, including reduced TNT formation, delayed fusion, altered multinucleation, and defective bone degradation. In vivo, deletion of either RIAM or Lpd in osteoclasts did not significantly affect bone mass, while disruption of the Rap1–Talin1 interaction slightly increased bone mass, consistent with an osteopetrotic phenotype. Together, these findings demonstrate that RIAM and Lpd exert distinct yet complementary functions and identify the direct Rap1–Talin1 interaction as a central mechanism controlling osteoclast fusion and bone resorption.

Talin1 is a key mediator of integrin activation and stabilisation through mechanical coupling to the actin cytoskeleton (Calderwood, Campbell and Critchley, 2013). In association with Kindlin3, it constitutes a core regulatory system essential for osteoclast function, as observed across multiple integrin pathways (Schmidt *et al*., 2011; Zou *et al*., 2013). Osteoclasts express numerous integrins with diverse ligand specificities, among which integrin αvβ3 plays a predominant role. Although several integrins act redundantly, particularly in bone degradation and sealing zone organisation (Schmidt *et al*., 2011), their individual contributions to osteoclast fusion remain insufficiently characterised. We show here, for the first time, that β1, β2, and β3 integrin families contribute, in a partially redundant manner, to osteoclast fusion. In particular, we observed that the suppression of β1 integrins unexpectedly increased the fusion capacity of osteoclast progenitors, a robust and reproducible observation. Given the central role of TNTs in facilitating osteoclast fusion (Dufrançais *et al*., 2025), it will be important to determine which integrins are present in these structures and how they contribute to TNT biogenesis and function, aspects that remain largely unexplored and warrant further investigation.

The functional redundancy of integrins during osteoclast fusion underscores the need to better define the upstream regulatory mechanisms. Upstream of Talin1, the small GTPase Rap1 acts as a signalling hub, integrating extracellular cues such as M-CSF and promoting the recruitment of Talin1 to integrins (Zou *et al*., 2013). Since the identification of RIAM (Lafuente *et al*., 2004) and Lpd (Krause *et al*., 2004) in 2004, their roles in integrin signalling and actin dynamics have been extensively studied. These two proteins bind to the same site on Talin1 and are therefore considered mutually exclusive (Lee *et al*., 2009, 2013), suggesting that they may act at distinct spatial or temporal stages during osteoclastogenesis. Consistent with this, we found that their inactivation produces different phenotypes in osteoclasts, associated with distinct subcellular localizations, although both proteins are found at actin-rich adhesion structures such as podosomes, where Talin 1 is also present. Despite belonging to the same MRL protein family, RIAM and Lpd exhibit divergent functional profiles in osteoclasts: Lpd appears to act during early fusion events, while RIAM functions mainly during later stages of bone resorption. These differences could reflect distinct regulatory mechanisms or downstream effectors, although the precise pathways remain to be elucidated. Their involvement is likely linked to a strong coupling between integrin activation and actin dynamics, which is consistent with previous findings (Lagarrigue *et al*., 2015). We also observed that the loss of Lpd, but not that of RIAM, reduces the formation of thick TNTs, which are essential for osteoclast fusion (Dufrançais *et al*., 2025). By analogy with its role in “sticky fingers” (Lagarrigue *et al*., 2015), structures involved in cell migration and sharing features with TNTs, Lpd could contribute to TNT elongation. Although TNTs are defined as long, non-adherent membrane extensions (>100 µm) (Zurzolo, 2021), integrins are likely necessary for their formation, particularly at the tips where anchoring to target cells occurs.

A better understanding of the mechanisms involved will require the generation of knockout mouse models for RIAM and Lpd, carrying mutations that selectively disrupt Talin1 binding or actin regulatory functions. Notably, Lpd differs from RIAM by its involvement in fast endophilin-mediated endocytosis, which regulates the internalisation of multiple membrane receptors (Vehlow *et al*., 2013; Boucrot *et al*., 2015). This process could be particularly relevant in osteoclasts, where it could control the distribution of fusogenic receptors such as DC-STAMP or key signalling receptors such as RANK, thereby directly influencing fusion capacity. Given its role in phagocytosis (Montaño-Rendón *et al*., 2022), Lpd could also control phagocytic cup-mediated cell fusion (Søe, Hobolt-Pedersen and Delaisse, 2015). Additionally, RIAM is primarily regulated by Rap1, with minor input from active Ras (Lafuente *et al*., 2004; Han *et al*., 2006; Lee *et al*., 2009), whereas Lpd preferentially interacts with other GTPases, notably Rac1 and M-Ras (Tasaka, Negishi and Oinuma, 2012; Law *et al*., 2013). These distinctions are important because Rac1 plays a well-established role in osteoclastogenesis (Korhonen *et al*., 2005; Vives *et al*., 2011), while the contribution of M-Ras remains less clearly defined. Together, these differences could explain the distinct functions of RIAM and Lpd, although further research is needed.

Beyond RIAM and Lpd, Rap1 can bind directly to the F0 and F1 domains of the Talin1 head, independently of these adaptors, which interact with the Talin1 rod domain (Vives *et al*., 2011). We found that selective disruption of these Rap1-binding sites produces a phenotype distinct from RIAM or Lpd deficiency, characterised by delayed rather than blocked osteoclastogenesis. This suggests that multiple Rap1–Talin1 interactions coexist, each contributing differentially to osteoclast function. An important question remains: do those interactions act in parallel or sequentially? One possibility is that direct binding of Rap1 to the head of Talin1 represents a fundamental constitutive recruitment mechanism of Talin1, to which a second interaction, RIAM or Lpd, would then be added to modulate the overall Rap1-Talin1 interaction. Osteoclasts are an ideal model for studying this question, as Talin1 participates in all stages of their differentiation and function. Data from other systems indicate that direct Rap1–Talin1 binding predominates in platelets, where RIAM and Lpd are minimally expressed (Zhu *et al*., 2017; Lagarrigue *et al*., 2018), while in T lymphocytes, RIAM can partially compensate for the disruption of this interaction (Lagarrigue *et al*., 2022). The situation in osteoclasts remains to be clarified.

Despite obvious in vitro defects, we show here that osteoclast dysfunctions are not fully reproduced in vivo in mice lacking individual effector pathways of Rap1 in myeloid lineages. In contrast, deletion of Rap1 or Talin1 (Zou *et al*., 2013), or loss of Kindlin3 (Schmidt *et al*., 2011), induces marked osteopetrotic phenotypes. The absence of such phenotypes in RIAM- or Lpd-deficient mice could reflect functional compensation between these proteins. Moreover, the use of LysM-Cre–mediated conditional knockouts may have limitations, as this system targets a broad myeloid lineage rather than specifically osteoclasts (Abram *et al*., 2014), and recombination efficiency may be incomplete in osteoclasts. Indeed, in vitro differentiation yields only 50–60% recombined osteoclasts (not shown), diluting observable phenotypes. Although we observed a high proportion (over 85%) of Ctsk-positive cells expressing Cre activity in vivo, the heterogeneity of osteoclasts remains poorly defined (Madel *et al*., 2019). The variability in integrin and actin cytoskeleton regulation within osteoclast subpopulations may exacerbate the complexity of observed phenotypes. The fact that disruption of the direct Rap1–Talin1 interaction induces a measurable phenotype in vivo, unlike deletion of RIAM or Lpd, suggests that this interaction could constitute a fundamental and ubiquitous mechanism for Talin1 recruitment. Osteoclasts also interact in complex and functional ways with other bone cells, particularly osteoblasts, suggesting that Rap1, Talin1, and Lpd may play additional roles in these cells. Although RIAM expression appears to be primarily restricted to hematopoietic cells, a potential role in osteoblasts cannot be ruled out. Future studies should incorporate alternative models, including global knockouts and more specific Cre drivers such as Ctsk-Cre, to better distinguish their respective roles in fusion and bone resorption. Furthermore, investigating these pathways in pathological contexts, such as osteoporosis, could reveal disease-specific regulatory mechanisms.

Osteoclasts, the main mediators of bone resorption, have long been considered key therapeutic targets in osteolytic diseases (Teitelbaum, 2000). Despite existing treatments, such as bisphosphonates or RANKL-neutralising antibodies, therapeutic options remain limited. From a translational perspective, actin-based adhesion structures, such as the sealing zone (Vives *et al*., 2015) and fusion-associated structures, represent subcellular features specific to osteoclasts. A better understanding of their regulation could therefore reveal new therapeutic targets. Although the regulatory mechanisms of integrins involving RIAM, Lpd, or Rap1–Talin1 are largely conserved and unlikely to be directly targeted, the identification of modulatory proteins within these pathways could offer new avenues for pharmacological intervention in osteolytic disorders.

In conclusion, this study highlights distinct yet complementary roles of RIAM and Lpd in osteoclast biology, emphasizing their different contributions to fusion and resorption activity. The findings further establish the direct Rap1–Talin1 interaction as a central and possibly common mechanism governing integrin activation in osteoclasts. The partial redundancy and complex interplay among integrins and their regulators underscore the need for more precise mechanistic models. Significantly, the discrepancies between in vitro and in vivo observations highlight the limitations of current experimental systems and the importance of more specific models. Ultimately, a better understanding of these pathways could lay the groundwork for the development of targeted therapies for osteolytic diseases.

## Acknowledgments

We acknowledge the TRI imaging and cytometry facilities, a member of the France-BioImaging national infrastructure, supported by the French National Research Agency (ANR-10-INBS-04), and in particular Eve Pitot, Pénélope Viana, and Emmanuelle Naser (IPBS). We also thank Mathias Krause, Anne Blangy and Isabelle Genero for fruitful discussions. We thank the Wellcome Sanger Institute for providing us with a Cas9 nuclease-expressing mouse line that they generated using the hCas9 plasmid obtained from Addgene.

This work was supported by the Centre National de la Recherche Scientifique (CNRS), Université de Toulouse, the Institut National de la Santé et de la Recherche Médicale, the Agence Nationale de la Recherche, the Fondation pour la Recherche Médicale (EQU202303016313). This study was supported by EUR CARe grant N°ANR-18-EURE-0003, within the framework of the Programme des Investissements d’Avenir, the ATIP program (CNRS INSB) and the Fondation ARC pour la recherche contre le Cancer, awarded to Frédéric Lagarrigue. Marianna Plozza, Camille Gorlt, Thibaut Sanchez, Ophélie Dufrançais, Camille Gorlt were supported by doctoral scholarships from Université de Toulouse, the Ligue Nationale contre le Cancer, the Fondation ARC pour la recherche contre le Cancer, the Fondation pour la Recherche Médicale and the EUR CARe graduate school. Rosa Nicole Freiberger holds a postdoctoral fellowship from the Fondation pour la Recherche Médicale.

## Supplemental figure legends

**Supplementary Figure 1** (related to Figure 1). Confocal microscopy images showing the localization of Talin1, RIAM and Lpd in human osteoclasts on glass slides, on day 8 of differentiation: F-actin (phalloidin, magenta), nuclei (DAPI, cyan) and Talin1/RIAM/Lpd (green). Scale bar: 10 µm.

**Supplementary Figure 2** (related to Figure 2). Experimental protocol for generating KO osteoclast progenitors (see the Methods section for details).

**Supplementary Figure 3** (related to Figures 2 and 3). **(A)** Flow cytometry analysis of β1, β2, and β3 integrin expression in murine osteoclasts on day 4 of differentiation. Representative histograms (n=3 experiments). **(B-C)** Role of β1, β2, and β3 integrins in osteoclast fusion. Microscopy analysis of osteoclast fusion osteoclasts on day 4 of differentiation. **(B)** Representative microscopy images: F-actin (phalloidin, white) and nuclei (DAPI, blue). Scale bar: 50 µm. **(C)** Quantification of the fusion index, n=3 independent experiments, 4 images/experiment, 1,000 nuclei/image, means ± SD are shown. **(D-E)** Role of β1, β2, and β3 integrins in bone degradation. Osteoclasts were cultured for 7 days on bone slices; after cell removal, the bones were stained with toluidine blue. **(D)** Representative images of bone degradation: eroded bone surfaces appear blue. Scale bar: 20 µm. **(E)** Quantification of eroded bone surface area (%) by semi-automatic quantification (n=3 independent experiments, 2 or 3 bone slices/condition, means ± SD are shown).

**Supplementary Figure 4** (related to Figures 4 and 5). Validation of the Talin1 R35E/R118E mutation in mice. **(A)** Schematic representation of conditional *Tln1* alleles before and after LysM-Cre–mediated recombination. In control (CTRL) mice, one allele carries the wild-type *Tln1* sequence and the other a floxed *Tln1* allele. In Talin1 RBM mice, one allele carries the R35E/R118E mutations, while the other is a floxed *Tln1* allele. In both genotypes, the Rosa locus contains a STOP cassette flanked by loxP sequences upstream of the eYFP reporter. During Cre-mediated recombination in myeloid cells, the floxed *Tln1* allele is excised along with the STOP cassette upstream of eYFP. In Talin1 RBM mice, recombination induces the expression of the mutant Talin1 R35E/R118E protein, while in CTRL mice, the wild-type Talin1 protein is expressed. In both cases the reporter gene eYFP is expressed after recombination. **(B)** Sanger sequencing confirms the presence of the R35E and R118E point mutations in the *Tln1* gene. Representative chromatograms show wild-type (WT) and Talin1 RBM sequences at the regions encompassing the R35E and R118E mutations (in blue). Nucleotide substitutions corresponding to amino acid changes (arginine to glutamate) are detected in the mutant samples, confirming the presence of the mutations altering Rap1 binding.

**Supplementary Figure 5** (related to Figures 4 and 5). Validation of the Talin1 R35E/R118E mutation by targeted proteomics and characterisation of osteoclast differentiation. **(A)** Schematic representation of the Talin1 protein region surrounding the R35E and R118E mutations. Tryptic peptides (pept1–3) generated by trypsin digestion and detectable by mass spectrometry in the wild-type protein are indicated. The positions of the R35E and R118E substitutions are highlighted in Talin1 RBM. **(B)** Mass spectrometry detection of Talin1-derived trypsin peptides corresponding to the regions shown in (A) in control (CTRL) and Talin1 RBM osteoclasts. **(C)** Heatmap showing the relative expression of selected macrophage- and osteoclast-associated genes in macrophages and osteoclasts. **(D)** Protein abundance (log_2_ intensity) of selected osteoclast-associated proteins and integrin regulators in CTRL and Talin1 RBM cells. **(E)** Protein abundance (log_2_ intensity) of selected integrin subunits in control (CTRL) and Talin1 RBM cells.

**Supplementary Figure 6** (related to Figure 6). X-ray µcomputed tomography analysis of distal cortical femora from wild-type mice and Lpd mKO, RIAM mKO, or Talin1 mRBM mice. Histograms represent the mean ± SD of trabecular thickness (left), trabecular number (middle) and cortical porosity (right). Each groups comprised 10 mice.

**Movie 1** (related to Figure 4). Time-lapse imaging of murine osteoclasts expressing wild-type Talin1 (CTRL) or Talin1 RBM on day 4 of differentiation. Images were acquired at 10-min intervals over a total imaging duration of 1 h and 40 min. Time is indicated as hh:mm. Scale bar, 20 µm.

**Movie 2** (related to Figure 6). 3D reconstruction of a confocal image stack showing eYFP expression in Cathepsin K-positive osteoclasts lining the femoral bone surface in control mice. Osteoclast identity was supported by multinucleation, elongated morphology, and bone-surface localization.

