## Supplemental Figures for "Uncoupling Osteoclast Fusion and Resorptive Activity Through Differential Talin1-Rap1 Signalling Pathways"

**Plozza et al. Supplemental Figure 1**

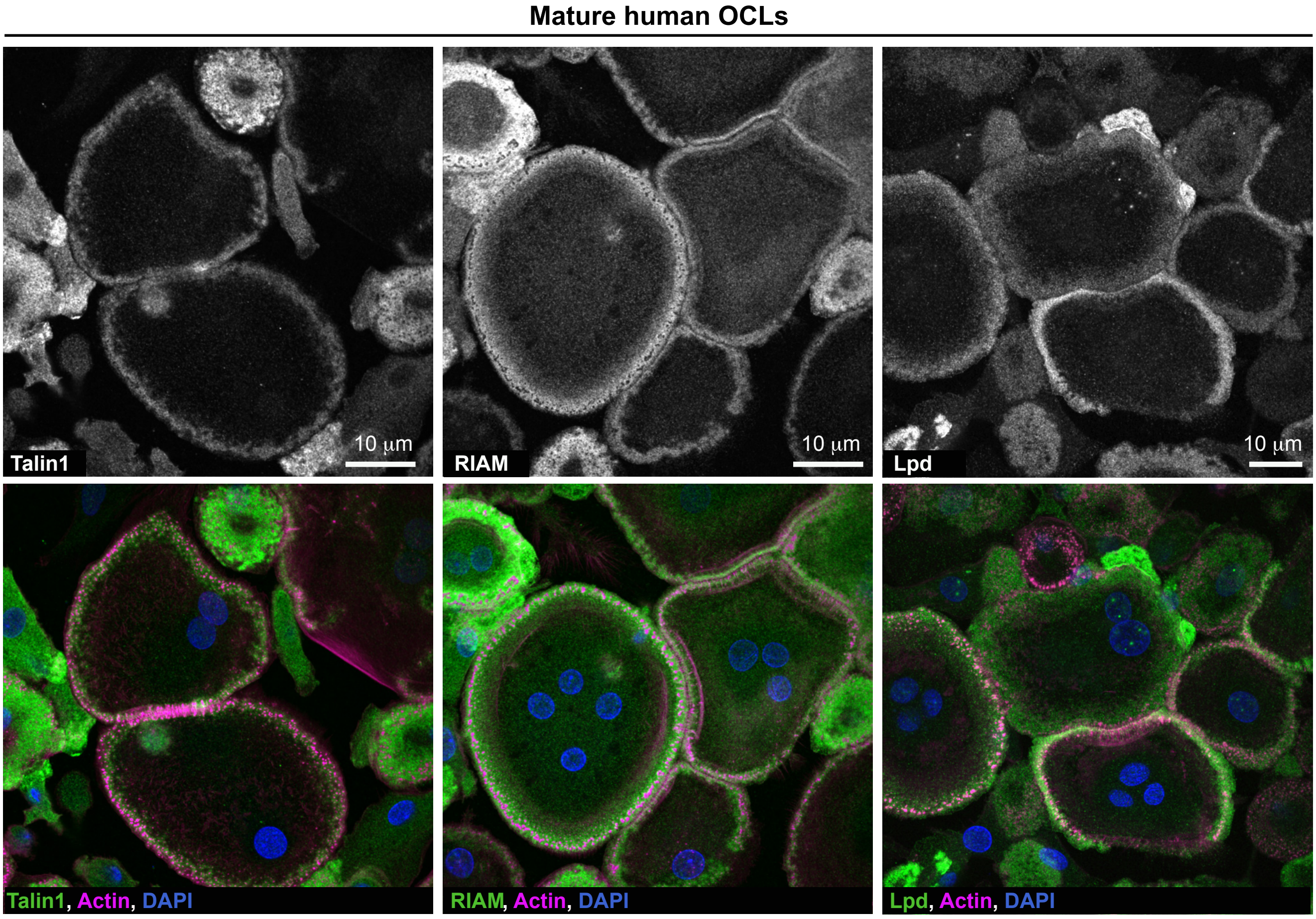

**Plozza et al. Supplemental Figure 2**

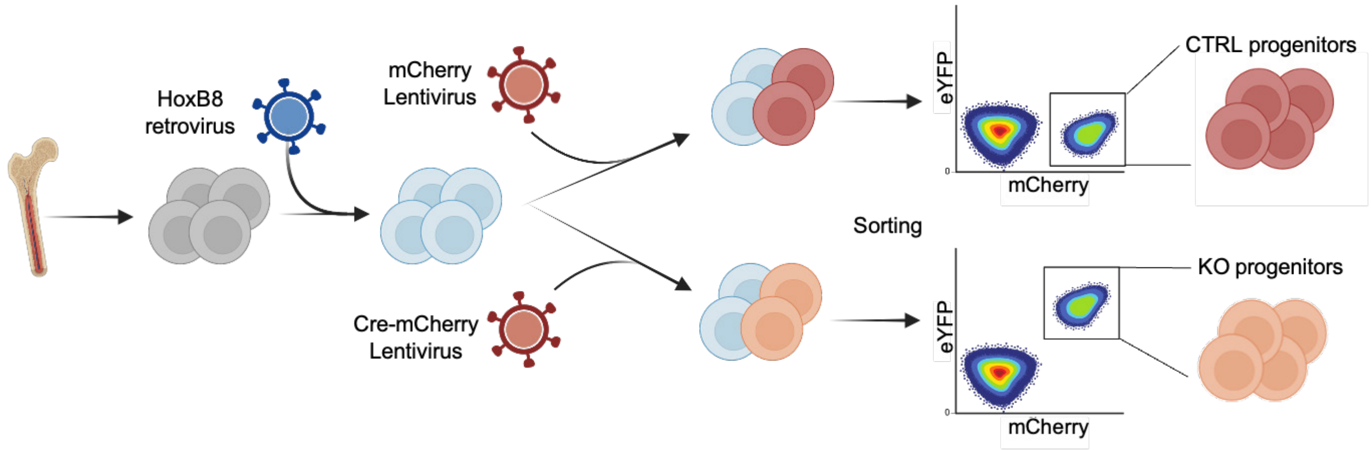

**Plozza et al. Supplemental Figure 3**

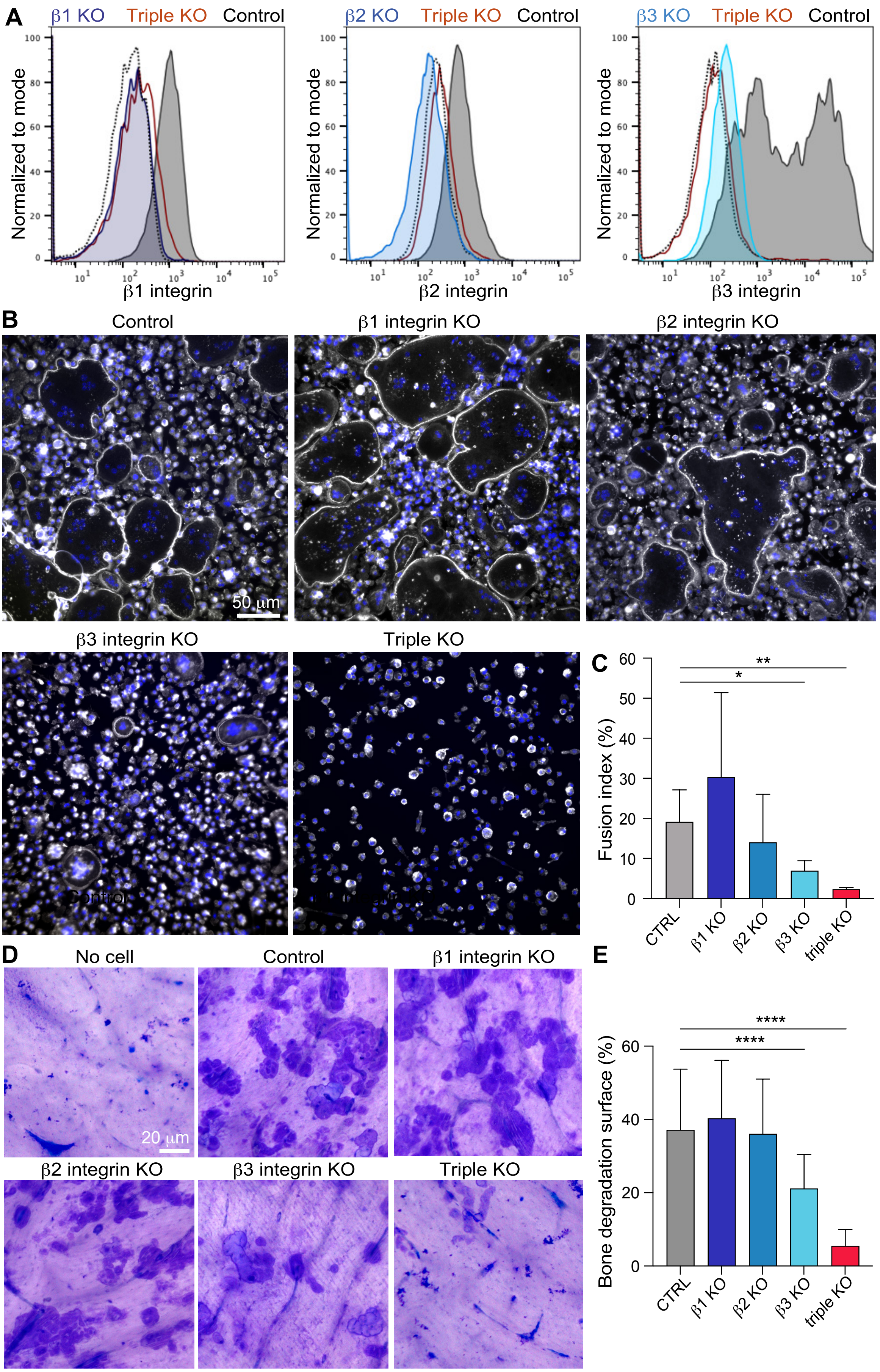

**Plozza et al. Supplemental Figure 4**

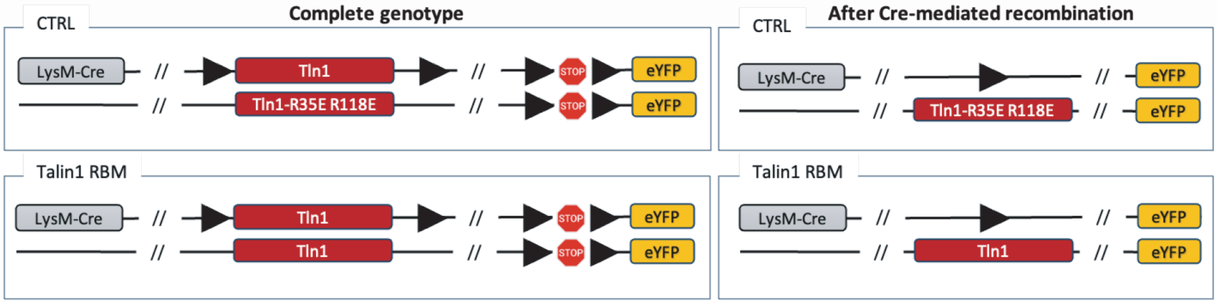

**Plozza et al. Supplemental Figure 5**

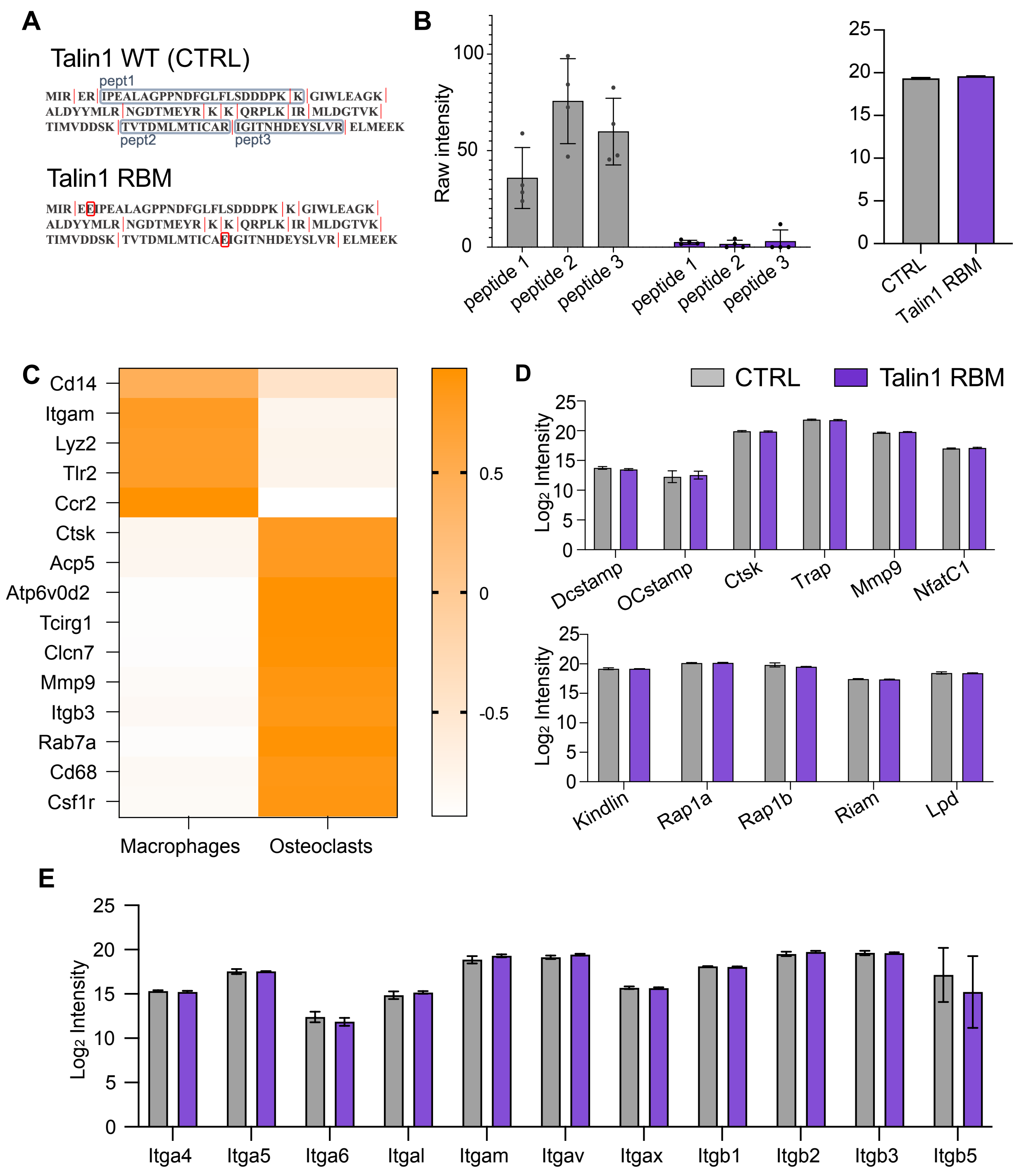

**Plozza et al. Supplemental Figure 6**

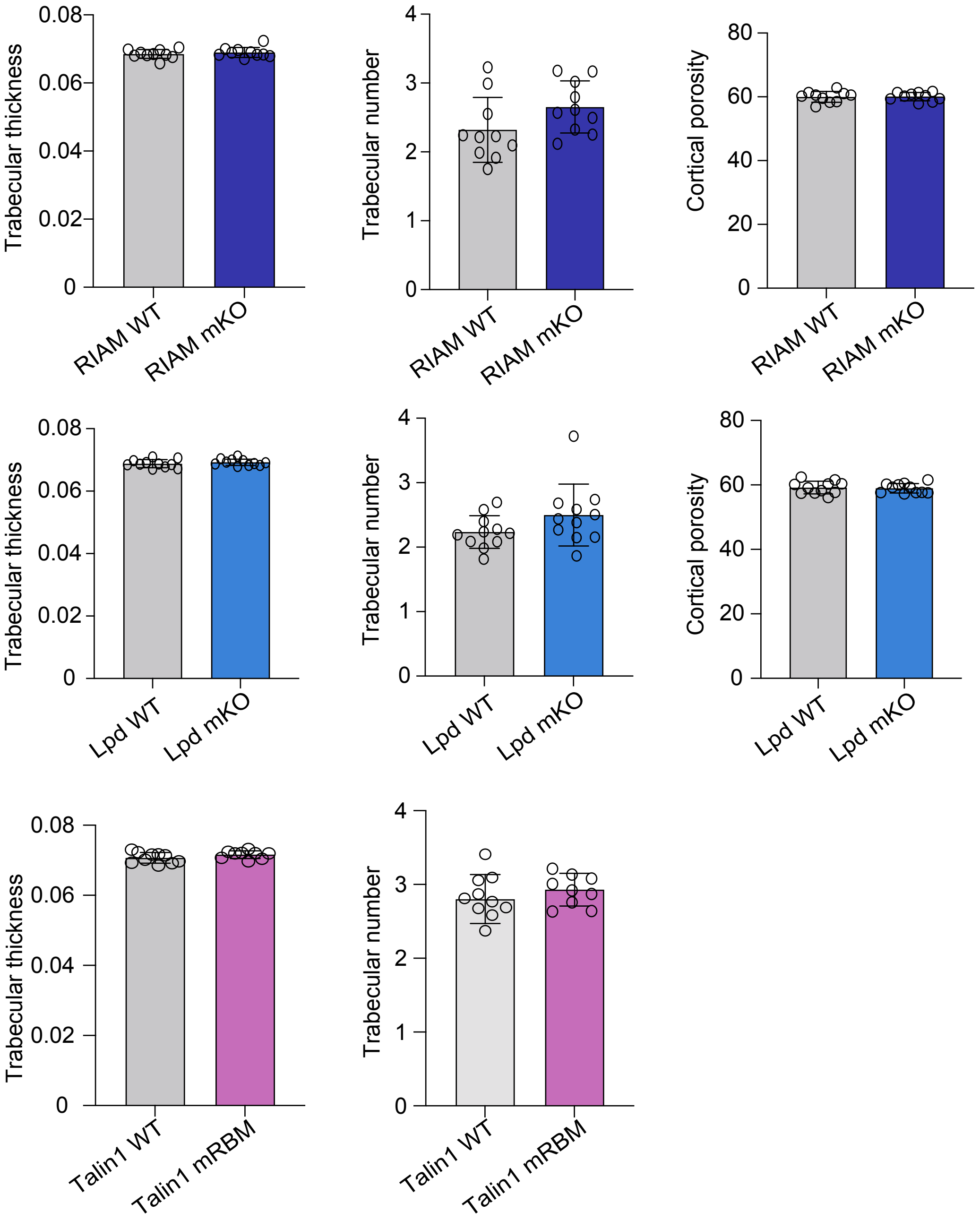
